# Smad1 and Smad5 differentially transduce BMP signaling during *in vitro* differentiation of mouse embryonic stem cells into dorsal interneurons

**DOI:** 10.64898/2026.06.30.735733

**Authors:** Salena Gallardo, Sandeep Gupta, Yahir Verdin, Cristian Rodriguez, Brian Chilin, Armo Derbarsegian, Gabriela Gajardo Del Real, Samantha J. Butler

## Abstract

A central unresolved question in development biology is how systems of overwhelming complexity arise from relatively few families of growth factors. Compounding this issue, signaling pathways often show signal convergence, where many ligands interact with fewer receptors, which then signal through a single second messenger complex. Here we investigate this question in the context of bone morphogenetic protein (BMP) signaling and its role directing dorsal spinal cord development, focusing on two receptor-regulated (R) Smads, Smad1 and Smad5. Multiple models have been proposed for their mode of action from acting redundantly through combined signal strength, to having distinct activities that drive different fate outcomes. We sought to distinguish between these models by generating CRISPR-edited *Smad1* and *Smad5* null mouse embryonic stem cell (ESC) lines to dissect the cell fate of activities of individual R-Smads, with a resolution not possible *in vivo.* Using a directed differentiation protocol for dorsal interneurons (dI), together with bioinformatic analyses, we have defined the roles of the R-Smads at key decision points along the dI specification timeline. Together, these findings support a model in which Smad1 and Smad5 play largely distinct roles in dorsal spinal cord development. While both R-Smads can activate canonical BMP-responsive transcriptional targets, they asymmetrically contribute to cell fate specification. Smad1 plays a restricted role, while Smad5 has a dominant role, regulating dorsal progenitor transcriptional dynamics and reiteratively directing the dorsal-most dI fates.

## Introduction

Bone morphogenetic proteins (BMPs) direct a remarkably wide array of cellular functions across different embryonic tissues, including proliferation, differentiation, migration, and cellular patterning (Hemmati-Brivanlou and Thomsen, 1995; Hogan, 1996; Kobayashi et al., 2005). Over 20 BMP ligands signal through ∼7 type I and type II serine/threonine kinase BMP receptors, which phosphorylate, and thereby activate, three canonical second messengers, the receptor-regulated (R) Smads (Smad1/5/8) (Fig. 1B) (Wang et al., 2014). The R-Smads complex with a co-Smad, Smad4, and translocate to the nucleus to regulate specific gene expression programs (Wrana et al., 1994; Zhang et al., 1996). The Smad complex acts as activators or repressors depending on the cofactors present and can also reshape chromatin structure, either directly or through the engagement of histone acetyltransferase (HAT) and histone deacetylase (HDAC) complexes (Akizu et al., 2010; Kim et al., 2011).

**Figure 1.**
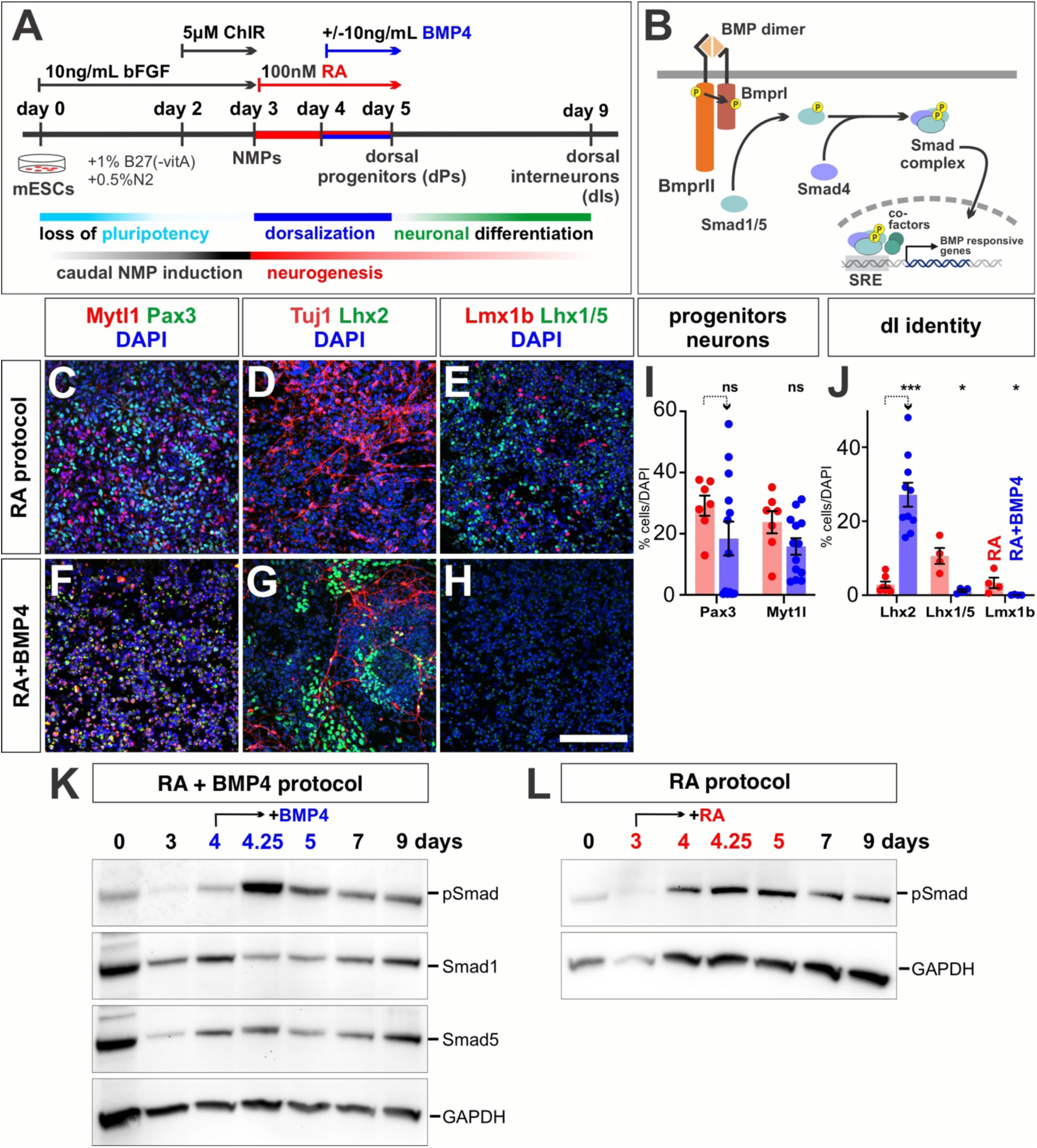
Timeline of Smad1/5 signaling in the mouse dI directed differentiation protocol (A) Schematic timeline of the 9-day RA±BMP4 *in vitro* directed differentiation protocols to derive dIs from mESCs. mESCs are first induced to lose pluripotency and enter the neuromesodermal progenitor (NMP) state. The addition of RA from day 3-5 directs NMPs towards a dorsal progenitor (dP) identity, which resolves into dI4s, dI5s and dI6s. Adding BMP4 concomitantly with RA from day 4-5, directs dPs towards dI1s, dI2s and dI3s. (B) Schematic of the canonical BMP signaling pathway. Upon BMP receptor binding, Smad1/5 are phosphorylated (p) and activated to serve as the key downstream mediators of BMP signaling in dorsal spinal cord development. (C-J) Day 9 *in vitro* cell cultures are effectively neutralized, containing both Pax3^+^ dPs and Myt1l^+^ post mitotic neurons (C, F, and I; RA: n=7 images, RA + BMP4: n=13). As previously demonstrated (Gupta et al., 2022; Rodriguez et al., 2026) The RA protocol induces the expression of Lhx1/5^+^ dI4s and Lmx1b^+^ dI5s (E, H and J; RA protocol n=7 (Lhx2) and n=4 (Lhx1/5, Lmx1b) RA + BMP4 protocol n=10 (Lhx2) and n=4 (Lhx1/5, Lmx1b), while the RA+BMP4 protocol induces the expression of Lhx2^+^ dI1s (D, G and J). (K, L) Western analyses assessed both Smad1 and Smad5 levels, and BMP signaling activity (p-Smad) across the timeline of the RA±BMP4 protocols. Smad1 and Smad5 are ubiquitously expressed throughout the RA + BMP4 protocol (K). BMP signaling, as measured by p-Smad immunoreactivity, is activated at consistent levels from day 4 onward in the RA protocol (L), but is markedly upregulated within 6 hours of recombinant BMP4 treatment in the RA+BMP4 protocol (K). Scale bar: 100µm Probability of similarity between control and experimental groups: * p<0.05, *** p<0.0005; two-way ANOVA

The ability of Smads to act as transcriptional regulators is well established. However, it remains unresolved how a single second messenger complex translates a diverse ligand input through a limited complement of receptors to result in the wide range of cellular outcomes directed by the BMPs. We are assessing this question in the dorsal spinal cord, where BMPs have reiterative roles directing cell fate (Andrews et al., 2019; Gupta and Butler, 2021) and axon guidance (Augsburger et al., 1999). The dorsal spinal cord contains six classes of dorsal interneurons (dIs), dI1-dI6, which are derived from six dorsal progenitor (dP) populations, dP1-dP6. These subtypes can be distinguished by their complement of transcription factors (Alaynick et al., 2011). BMP signaling is known to specify the dorsal-most spinal cord, i.e., the roof plate (RP), dI1, dI2 and dI3 (Andrews et al., 2017; Lee et al., 2000; Lee et al., 1998; Wine-Lee et al., 2004). The intermediate spinal cord, i.e. dI4-dI6, is thought to be specified by retinoic acid (RA) (Gupta et al., 2022). RA is present in the surrounding paraxial mesoderm and has an earlier role directing neuralization (Diez del Corral et al., 2003; Novitch et al., 2003).

Previous studies have suggested that R-Smads function interchangeably during embryogenesis (Arnold et al., 2006). However, R-Smads are found in different compartments of the developing spinal cord: Smad5 is present in highest levels in progenitors, whereas Smad1 is enriched in post-mitotic neurons and their processes, and Smad8 is completely absent (Hazen et al., 2012). This distribution pattern is consistent with Smad1 and Smad5 having distinct, rather than redundant, roles in spinal cord development. Our loss-of-function conditional genetic approaches in mouse and overexpression studies in chicken supported this hypothesis, suggesting that Smad5 mediates the cell fate specification activity of the BMPs, while Smad1 regulates axon outgrowth (Hazen et al., 2012). However, other studies have differed in their conclusions; RNA inhibition approaches in chicken embryos suggested that Smad1 and Smad5 redundantly mediate dI fate specification (Le Dreau et al., 2012). This finding led to the signal strength model which proposes that the level of combined Smad1/5 activity, rather the identity of individual Smads, determines cell fate (Le Dreau et al., 2014). R-Smads are phosphorylated (p) in a gradient in the spinal cord (Hazen et al., 2012), consistent with the signal strength model. However, we subsequently found that different BMPs can direct similar levels of pSmad1/5, yet result in markedly different fate outcomes (Andrews et al., 2017), suggesting that R-Smad identity, rather than signal strength, is the key determinant of cell fate.

*In vivo* approaches have inherent limitations. Acutely misexpressing genes in chicken is performed in a background of reiterative endogenous signaling, while RNA inhibition approaches can be non-specific, targeting multiple related genes. In contrast, the cre-lines used in conditional knockout approaches in mice may lack tissue specificity, or have perduring gene function that masks phenotypes especially if protein turnover is slow. To overcome these limitations, we have developed a new mouse embryonic stem cell (ESC) model system where CRISPR/Cas9 genome editing (Jiang and Doudna, 2017) was used to generate lines lacking either Smad1 (Smad1Δ) or Smad5 (Smad5Δ). Stem cell models have been powerful systems for interrogating complex developmental patterning and transcriptional mechanisms (Gouti et al., 2014; Panchision et al., 2001; Sagner et al., 2018), including ventral spinal cord development (Mazzoni et al., 2013; Sagner et al., 2018). They generate large populations of cells relatively synchronously executing the same developmental decisions and permit experimental control over signaling inputs in ways that are not feasible *in vivo*.

Here, we have used established directed differentiation protocols for dIs (Gupta et al., 2022; Rodriguez et al., 2026) as a platform to dissect the individual roles of Smad1 and Smad5 in dorsal spinal cord development. These protocols generate dIs via a neuromesodermal progenitor (NMP) intermediate (Gouti et al., 2014), and thereby recapitulate the endogenous program of dorsal spinal cord development (Gupta et al., 2025; Gupta et al., 2022). The RA protocol is enriched for intermediate dIs (dI4-dI6), while the RA+BMP4 protocol most effectively directs the dorsal-most dIs (dI1-dI3) (Gupta et al., 2022). By applying bioinformatic approaches at key decision points along the dI specification timeline in control, Smad1Δ and Smad5Δ mESC lines, we have defined the individual contributions of each R-Smad to dorsal spinal cord development. We find that while both R-Smads activate canonical BMP-responsive transcriptional targets, their contributions to cell fate decisions are asymmetric. Smad1 plays a restricted role suppressing specific dorsal fates, while Smad5 is the dominant second messenger, directing transcriptional dynamics across the dorsal progenitor domain (dP1-dP6) and reiteratively directing the dorsal-most dI fates. Together, these findings support a model in which Smad identity, rather than signal strength alone, determines the diverse cell fate outputs of BMP signaling in the dorsal spinal cord.

## Materials and Methods

### Differentiation of mouse embryonic stem cells into dIs

Control, Smad1Δ, and Smad5Δ mESCs were differentiated into dIs using two-dimensional adherent protocol as previously described (Rodriguez et al., 2026).

### Western analysis

Cells were washed with 1x PBS and cold Pierce® RIPA buffer (Thermo Scientific) supplemented with a 1x protease inhibitor cocktail (Roche) and 1x phosphatase inhibitor cocktail (Roche) was directly added to multiple wells of a 24 well differentiation plate. The lysed cell suspensions were pooled, held on ice for 15 minutes, then centrifuged while at 4°C for 15 mins. at 14,000 x g. A BCA assay was performed on the supernatant to determine protein concentrations. 40ug of protein was used as input for SDS-PAGE, using a gradient 4-12% Bis-Tris SDS gel (GenScript) followed by transfer onto PVDF membranes (Millipore Sigma). The membranes were blocked using 5% non-fat dry milk for 1 hour (Bio-Rad Laboratories). The blocked membranes were then incubated with the primary antibodies (key resources table) at 4°C overnight. Thereafter, the membranes were washed three times with TBST (20 mM Tris, 150 mM NaCl, 0.1% Tween 20), incubated with species-specific horseradish peroxidase-conjugated secondary antibodies (Jackson ImmunoResearch Laboratories) for an hour at room temperature and then washed twice with TBST and once with TBS. The chemiluminescent bands were analyzed using the Pierce Femto Chemiluminescence kit on the ChemiDoc^TM^ imaging system (Bio-Rad). If the Western blots were reused (Fig. 1K), the membrane was stripped using the RestoreTM PLUS buffer (Thermo Scientific) and re-analyzed to confirm the absence of protein signal.

### Immunohistochemistry

Cell cultures were washed with 1x PBS and fixed with fresh cold 4% PFA for 8 min in the well. Following fixation, cultures were washed twice with 1xPBS to remove any remaining PFA. Cells were blocked with 1xPBS with 1% heat inactivated horse serum for 1 h and primary antibodies are added in the blocking solution for an overnight incubation at 4°C. Following washes, species appropriate secondary antibodies (Jackson Immuno Research Laboratories) were added in 1x PBST (PBS + 0.1% Triton 20) for 1hr. The cultures were then washed with 1x PBST to remove traces of secondary antibodies and counterstained with DAPI to stain nuclei. Plates were then imaged on Zeiss LSM800 inverted confocal microscope at 20x magnification.

### Generation of Smad1 and Smad5 deficient (Δ) mESC lines

A CRISPR-mediated genomic deletion strategy was utilized as described (Ran et al., 2013), to generate Smad1 and Smad5 deficient (Δ) mESC lines. Briefly, guide RNAs (gRNAs) targeting upstream and downstream of exon2 in either the *Smad1* or *Smad5* gene locus were cloned into the pSpCas9(BB)-2A-Puro V2.0 plasmid. Various combinations of gRNA constructs were then delivered to the cell via the Lonza 4D-Nucleofector® X Unit. Nucleofected cells were plated at a single cell density onto tissue culture plates containing mESC media supplemented with puromycin. Colonies that grew in the presence of puromycin were isolated, expanded, and further characterized to confirm the correct genomic deletion.

### Characterization of Smad1Δ and Smad5Δ mESC lines

We confirmed successful excision of the exon 2 locus by PCR, Sanger sequencing, and western blotting. Candidate Smad1Δ and Smad5Δ mESC lines were grown on 0.1% gelatin-coated plates and passaged at 75% confluency. After two passages, cells were washed with 1x PBS and incubated with 0.25% trypsin at 37°C for 3 min. Trypsin dissociation activity was neutralized with mESC media in a 1:1 ratio and cells were collected and centrifuged at 1,000 RPM for 5 min. Cell pellets were resuspended in 1x PBS and evenly partitioned for downstream analysis.

#### PCR and Sanger sequencing

Genomic DNA was isolated from the resuspended cell pellet using the QIAamp® DNA Mini kit and following the manufacturer’s protocol. PCR amplification across the CRISPR-targeted exon 2 genomic region was performed using primers designed to anneal outside the deleted interval. PCR conditions were: 95 °C for 2 min; 30 cycles of 95 °C for 15 s, 55.7 °C for 30 s, and 72 °C for 3 min; followed by 72 °C for 5 min and held at 4 °C. Amplicons were resolved on either a 1.25% (Smad1Δ) or 2% (Smad5Δ) agarose gel. After visually confirming the presence of a large genomic deletion, the remaining PCR products were cleaned up using the MinElute^TM^ PCR purification Kit (Qiagen) and submitted for Sanger sequencing. PCR amplification of genomic DNA from the SF20 mESC line resulted in two different sized PCR products; each band was gel extracted using the Monarch® DNA Gel Extraction Kit (New England Biolabs) and submitted separately for Sanger sequencing. Sequence traces were aligned to the GRCm38 *Mus musculus* genome using the ApE (A Plasmid Editor) program to determine the size and boundaries of the genomic deletion.

#### Western Analysis

Cell suspension was centrifuged for 5 min. at 1000 RPM and resuspended in 1x Pierce® RIPA buffer (Thermo Scientific) supplemented with a 1x protease inhibitor cocktail (Roche). The lysed protein suspension was held on ice for 15 mins. then centrifuged while at 4°C for 15 mins. at 14,000 x g. The supernatant was subjected to SDS-PAGE and the remaining steps were carried out as previously described in the above methods section.

### Bulk-RNA sequencing and data processing

#### Sample collection, library preparation, and sequencing

Total cell lysate was obtained in buffer RLT (Qiagen) at different time points from three independent differentiations (biological replicates) conducted in parallel, for each genotype: control (MM13), Smad1Δ (C2, C34, C36) and Smad5Δ (SF20, SF27, SF43). RNA extraction was performed using RNeasy mini kit (Qiagen) and the quality was determined using Agilent Technologies 2100 Bioanalyzer. Stranded libraries were constructed using Universal plus mRNA sequencing kit (NuGEN) and sequenced onto 1.5 lanes of NovaseqS4 to generate a minimum of 40 million reads/sample (with the exception of day 4-C34 at 24M reads). Notably, 4 samples failed library prep (day 0-SF20, day 0-SF27, day 3-SF43, day 9-B4-ControlA) and 3 samples did not generate an adequate library for sequencing (day 3-SF27, day 5-RA-SF27,day 9-RA-SF20). A fourth differentiation was performed to replace the day 0 through day 5 samples, resulting in a total of 79 samples that were sequenced.

#### Data processing

Raw sequencing reads were aligned to the *Mus musculus* GRCm38 reference genome using the STAR aligner and gene-level counts were generated with featureCounts (Subread package). Genes with fewer than 10 total reads across all samples were removed prior to normalization. Quality control was performed using Pearson correlation and principal component analysis (PCA) to assess replicate similarity and genotype clustering. Differential expression analysis was conducted using the DESeq2 package in R. Differentially expressed genes were extracted using DESeq2 contrast statements corresponding to: (1) time effects within WT, Smad1Δ, and Smad5Δ, and (2) genotype differences (Smad1Δ vs WT and Smad5Δ vs WT) at day 0, day 3, and day 4. For the later differentiation stages, samples collected at day 4, day 4.25, day 5, and day 9 under RA or RA+BMP4 media conditions were analyzed using a grouped DESeq2 design. Differential expression analyses were performed to compare RA vs RA + BMP4 at matched timepoints (day 4.25 and day 5) within each genotype. Significant genes were defined as those with an adjusted p-value (padj) < 0.05 and absolute log2FoldChange >1. Venn-like overlap analyses were performed to identify genes uniquely or commonly regulated across WT, Smad1Δ, and Smad5Δ backgrounds at day 4.25 and day 5. Notably, 3 samples were excluded due to a file level read error encountered during FASTQ data retrieval (day 0-ControlB, day 4.25-B4-ControlC, day 9-RA-C34).

#### Gene Ontology (GO) enrichment analysis

Gene Ontology enrichment analysis was performed using the clusterProfiler package. Biological process enrichment was tested using the Benjamini–Hochberg method for multiple-testing correction, with significance thresholds of p-adjusted < 0.05 and q-value < 0.2.

#### Generation of heatmaps

Variance-stabilized expression values were averaged within each timepoint–genotype–media group, row-centered by Z-scaling, and visualized as heatmaps using the pheatmap package in R.

### ATAC-sequencing

We performed parallel differentiations of three control (biological replicates) mESC lines and collected samples for ATAC sequencing at day 0, D3, day 4, day 4.25 (6 hours ± the addition of BMP4) and day 5 (24 hours ± the addition of BMP4).

#### Sample preparation and sequencing

Samples were prepared as previously described (Buenrostro et al., 2015). Cultures were dissociated with 0.25% cold trypsin for 5 min. Trypsin activity was stopped with the addition of trypsin inhibitor (Sigma-Aldrich) in 1:1 ratio. Cells were then pelleted by centrifugation at 1000RPM and resuspended in 1x PBS. Cells were counted and 75,000 cells were collected and centrifuged at 500 x g for 5 mins at 4°C. The cell pellet was resuspended in 50 uL of cold lysis buffer (10mM Tris-Cl pH7.4, 10mM NaCl, 3mM MgCl₂, 0.1% IGEPAL CA-630) and immediately centrifuged at 500 x g for 10 min at 4°C. Nuclei were resuspended in transposition reaction mix containing TDE1 Tn5 Transposase and 1x TD reaction buffer (Illumina) and incubated at 37°C for 30 min. Transposed DNA was purified using the MinElute™ PCR Purification Kit (Qiagen) per the manufacturer’s instructions. Libraries were sequenced on a NovaSeq S4 (XP workflow, paired-end 2×100 bp), yielding an average of ∼120 million reads per sample.

#### Preprocessing

Nextera adapter sequences were trimmed and low-quality bases removed using Trimmomatic. Trimmed reads were aligned to the GRCm39 mouse reference genome using Bowtie2. Resulting SAM files were converted to BAM format, coordinate-sorted, and indexed using SAMtools. PCR duplicates were identified and removed using Picard MarkDuplicates. Reads mapping to ENCODE blacklisted regions were excluded using BEDtools intersect with the mm39 blacklist (v2). Filtered BAM files were merged across biological replicates per condition, re-sorted, and re-indexed using SAMtools. Open chromatin regions were identified by peak calling with MACS3. Normalized coverage tracks were generated using deeptools bamCoverage with counts per million (CPM) normalization and a bin size of 25 bp.

#### Unified Peak Set and ATAC Signal Quantification

A unified set of regulatory regions was generated by centering fixed-width windows on peak summits identified across all conditions, resulting in a unified peak set of 141,592 peaks. Genomic annotation of the unified peak set was performed using annotatePeaks.pl from the HOMER suite against the mm39 reference genome. ATAC signal over the unified peaks was quantified using computeMatrix (deepTools) in reference-point mode, anchored at the center of each peak with 500 bp flanking regions on each side. Chromatin accessibility was visualized using plotHeatmap and plotProfile from deepTools, displaying mean CPM signal centered on peak summits across samples.

#### Motif enrichment analysis

Treatment-responsive regulatory elements at 24 hours were defined as unified ATAC-seq peak regions exhibiting a log₂ fold-change > 0.5 in normalized accessibility signal between treated (+BMP4) and untreated (RA) cells. Transcription factor motif enrichment analysis was performed on these peaks using HOMER (findMotifsGenome.pl) against the mm39 reference genome.

#### Promoter-Centric Analysis

Promoter regions were defined as ±2 kb windows centered on the transcription start site (TSS) of each annotated gene. BMP4 responsive ATAC peaks at 24 hours were intersected with promoter regions using BEDtools intersect, identifying 16 promoter regions with increased chromatin accessibility. These regions were mapped to their nearest gene and intersected with bulk RNA-seq data from the same timepoint comparison (day 5 ± BMP4). Genome browser tracks displaying CPM-normalized ATAC signal at the promoters of candidate genes (*Id2*, *Msx1*, and *Wfikkn1*) were visualized over a ∼±2 kb window centered on the annotated TSS of each gene.

### Single-cell RNA sequencing

We performed parallel differentiations of control (MM13), Smad1Δ (C36) and Smad5Δ (SF43) mESC lines and collected samples for single-cell sequencing at day 9 of the dI directed differentiation protocol.

#### Preparation of the single cell suspension

Control, Smad1Δ, and Smad5Δ day 9 cultures, from both RA and RA + BMP4 protocols, were dissociated with 0.25% cold trypsin for 5 min. Trypsin activity was stopped with the addition of trypsin inhibitor (Sigma-Aldrich) in 1:1 ratio. Cells were then pelleted by centrifugation at 1000RPM and resuspended in 1x PBS. Dead cells were removed using MACS dead cell removal kit (Miltenyi Biotec Inc.) by following the manufacturer’s protocol. Eluted live cells were then suspended in 1X PBS containing 0.04% BSA solution (400μg/ml) for the library preparation.

#### Library preparation and sequencing

∼10,000 live cells/conditions were used to construct single-cell specific cDNA libraries using protocol described in 10x Genomics chromium single cell 3′ reagent kit (v3.1 Chemistry). Briefly, cells were partitioned into nanoliter-scale Gel-Beads-in-emulsion (GEM) using the 10x Chromium controller. Each GEM contains a unique barcode which is shared among the cDNA generated from a single cell. Cells were then lysed, and cDNA synthesis and feature barcoding were performed in the GEMs. The sequencing libraries were recovered using Magnetic separation and the quantity and quality of cDNA were assessed by Agilent 2100 expert High Sensitivity DNA Assay. Libraries were sequenced across two lanes of an Illumina NovaSeq X Plus using a 10B flow cell.

#### Preprocessing and filtering

Raw 10x Genomics count matrices were imported into Seurat. For each sample, cells were filtered based on thresholds determined from QC plots. Specifically, cells were retained if they met sample-specific criteria for gene complexity (nFeature_RNA > 250, with upper limits of 7,300–8,000 depending on sample), total UMI counts (800 < nCount_RNA < 38,000–50,000), and mitochondrial content (percent.mt < 10%). These thresholds were applied independently to each genotype–condition dataset (WT, Smad1Δ, and Smad5Δ under RA or BMP4 conditions) to remove low-quality cells, empty droplets, and potential doublets. Cells failing to meet any of these criteria were excluded prior to library-size normalization, dimensionality reduction, and subsequent downstream analysis.

#### Data integration

Following per-sample quality control, the six filtered Seurat objects (control, Smad1Δ, and Smad5Δ under BMP4 and RA conditions) were merged into a single Seurat object and split by genotype–treatment group using SplitObject. Each sub-object was independently normalized with SCTransform, and the top 3,000 variable features were selected across all groups using SelectIntegrationFeatures and prepared for integration with PrepSCTIntegration. Cross-dataset integration anchors were identified with FindIntegrationAnchor and used to generate a single batch-corrected object via IntegrateData, both using SCT normalization.

#### Dimensionality reduction, clustering and Cell Type annotation

Following integration, principal component analysis (PCA) was performed on the corrected integrated assay using RunPCA. Uniform Manifold Approximation and Projection (UMAP) was computed using the top 30 principal components as input via RunUMAP. Unsupervised clustering was performed using a shared nearest-neighbor graph constructed with FindNeighbors across the same 30 PCs, followed by Louvain community detection at a resolution of 0.8 using FindClusters.

Cell type identities were assigned to each Seurat cluster using the ScType automated annotation framework (Ianevski et al., 2022). ScType was run on the SCT-normalized, scaled assay with a custom major lineage marker gene file. Cluster-level annotations generated by ScType were used to label cells in a final UMAP visualization, in which each cell was colored by its ScType-assigned major lineage identity. To visualize and quantify the proportion of NMP derivatives, the progenitor, roof plate, dorsal progenitor, and neuron clusters were grouped into a single "neural" category, while mesoderm and vascular clusters were grouped into a single "mesoderm" category. To enable fair visual comparison of cell type composition across samples, cells were down sampled prior to plotting. The minimum cell counts across all genotype–treatment groups was determined, and each group was randomly subsampled to this number using a fixed random seed (seed = 123) to ensure reproducibility. A down sampled Seurat object was generated from the retained cells and used exclusively for visualization (Fig.5D, 5E, 5G, 5H).

#### Cell type proportion analysis

Cell type proportions were quantified per genotype–treatment group by tabulating the number of cells assigned to each cluster within each sample. Proportions were expressed as a percentage of total cells per group and visualized as pie charts for each of the six genotype–treatment combinations. Category labels were displayed within each slice only when the proportion exceeded 5% of total cells.

#### Marker gene dot plot

To validate ScType cluster annotations, a dot plot was generated using a curated panel of known marker genes across all annotated cell type clusters. Expression was visualized from the SCT-normalized assay using Seurat’s DotPlot function. Dot size reflects the percentage of cells within each cluster expressing a given gene, and dot color reflects the scaled mean expression level. Genes absent from the dataset were excluded prior to plotting.

#### dP and dI subset, reclustering, and annotation

Cells annotated as dorsal progenitor or neuron in the full integrated object were extracted as a subset. PCA, UMAP, and unsupervised clustering were performed as previously described. Cell type annotation of the filtered neural subset was performed using ScType with a curated dP and dI marker panel. Upon inspection of dorsal progenitor marker gene expression, the dP3 population was not captured as a discrete cluster. To recover this population, cells initially annotated as dP2 that expressed *Ascl1* above a normalized expression threshold of 0.25 were manually relabeled as dP3. This revised annotation was stored as a separate metadata field and used for all downstream visualization and analyses of the dP/dI subset. Notably, prior to clustering, an anomalous neural cluster was identified upon inspection and its relative contribution to each genotype–treatment group was quantified. As this cluster represented a minor and inconsistent population, it was excluded from all subsequent analyses.

Down sampled UMAPS were generated as previously described (Fig. 6D,E,G,H). Cell type proportion analysis was performed as previously described. Marker gene dot plot was generated as previously described.

### RNA velocity analysis

#### Data preparation and preprocessing

Spliced and unspliced RNA count matrices were generated for each sample using velocyto (La Manno et al., 2018) and loaded into Python using the AnnData library. Each sample was then filtered to retain only cells present in the complete neural atlas (Fig. 6A). The six filtered samples were concatenated into a single AnnData object. To maintain consistency with the Seurat analysis, UMAP coordinates and cluster annotations were transferred directly from the neural atlas Seurat object by loading the exported embeddings and cluster label metadata and mapping them to the AnnData object. Preprocessing for velocity estimation was performed using scVelo. Genes were filtered and normalized using a minimum shared spliced/unspliced count threshold of 20 and a minimum total count of 20. A nearest-neighbor graph was constructed using 30 neighbors, and first- and second-order moments of the spliced and unspliced count distributions were computed across the neighborhood graph in preparation for velocity estimation.

#### Analysis of integrated neural atlas

RNA velocity was estimated using the stochastic model implemented in scVelo. The spliced/unspliced read proportions were inspected per cluster to confirm adequate unspliced RNA recovery across cell types. Velocity vectors were projected onto the UMAP embedding as a streamline plot, with cells colored by cluster identity, to visualize the directionality and continuity of transcriptional trajectories across the dP/dI populations.

#### Identification of driver genes in control cells

To characterize baseline transcriptional dynamics, velocity length, confidence, and confidence transition were assessed in control cells stemming from both RA and RA+BMP4 protocols. Velocity genes were ranked across clusters in control cells using the scv.tl.rank_velocity_genes function in velocyto, with cells grouped by cluster identity and a minimum correlation threshold of 0.3. Phase portraits were generated for selected top velocity genes including *Magi1* and *Smoc1*.

#### Velocity analysis in Smad1Δ and Smad5Δ data sets

Velocity streamline plots were generated for each of the six groups (Control, Smad1Δ, and Smad5Δ under RA and RA + BMP4 protocols). Velocity magnitude was quantified across groups and clusters by computing the mean velocity length, velocity confidence, and velocity confidence transition per group–cluster combination.

dI1 cells from RA+BMP4 treated samples were isolated and the mean absolute velocity magnitude was computed per gene for each group. Genes with negligible velocity signal across all groups were excluded. For each remaining gene, pairwise comparisons between control and each mutant were performed using the Wilcoxon rank-sum test, and p-values were corrected for multiple testing using the Benjamini–Hochberg false discovery rate method. Candidate genes with significantly reduced velocity in mutant relative to control cells (FDR < 0.05, delta > 0) were identified separately for Smad1Δ and Smad5Δ comparisons and ranked by effect size. *Nav3* and *Cadm2* were identified as genes with significantly reduced mean velocity magnitude in Smad1Δ and Smad5Δ cells relative to control. The normalized expression of *Nav3* and *Cadm2* was visualized on the dI1 UMAP for each RA+BMP4 treated group.

### Quantification and statistical analysis

#### Image quantification

To count the number of nuclei, images were converted to 8bit image in ImageJ, their threshold intensity was adjusted to capture only the fluorescent nuclei, and the number of nuclei counted using the analyze particle tool in ImageJ.

#### Statistics

Data are represented as mean ± SEM (standard error of the mean). Tests for statistical significance were performed using Prism software. Values of p < 0.05 were considered significant in all cases.

## Key Resource Table

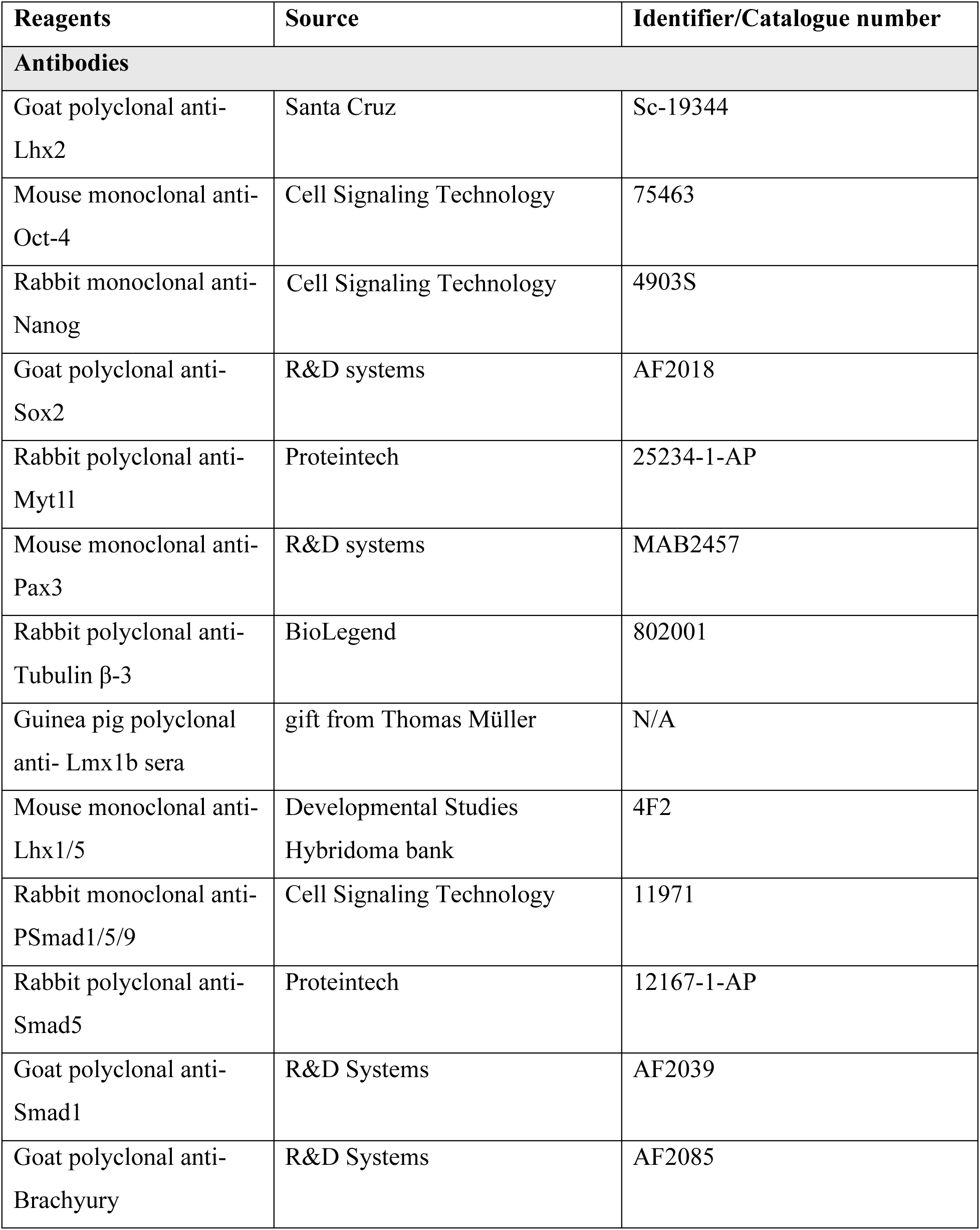
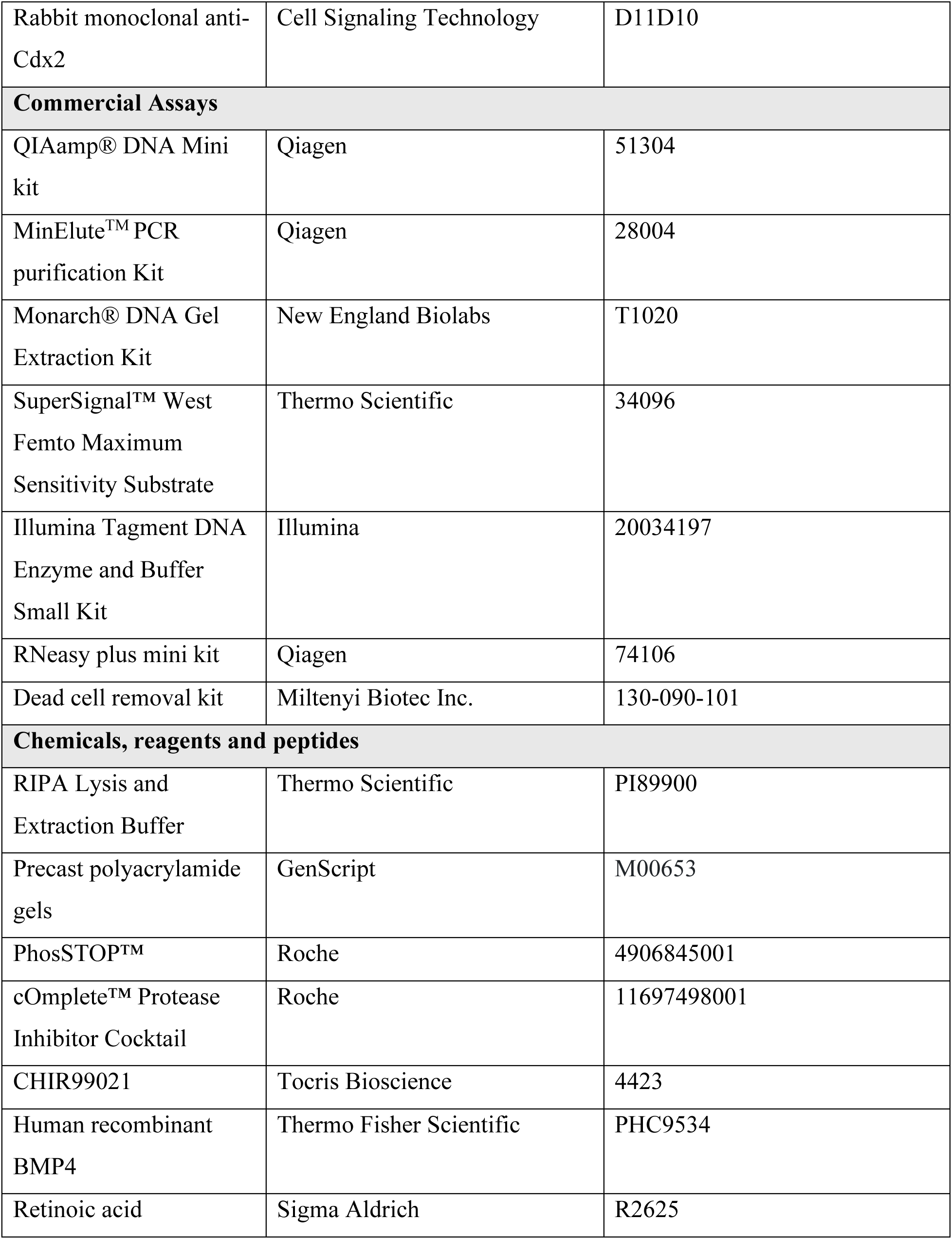
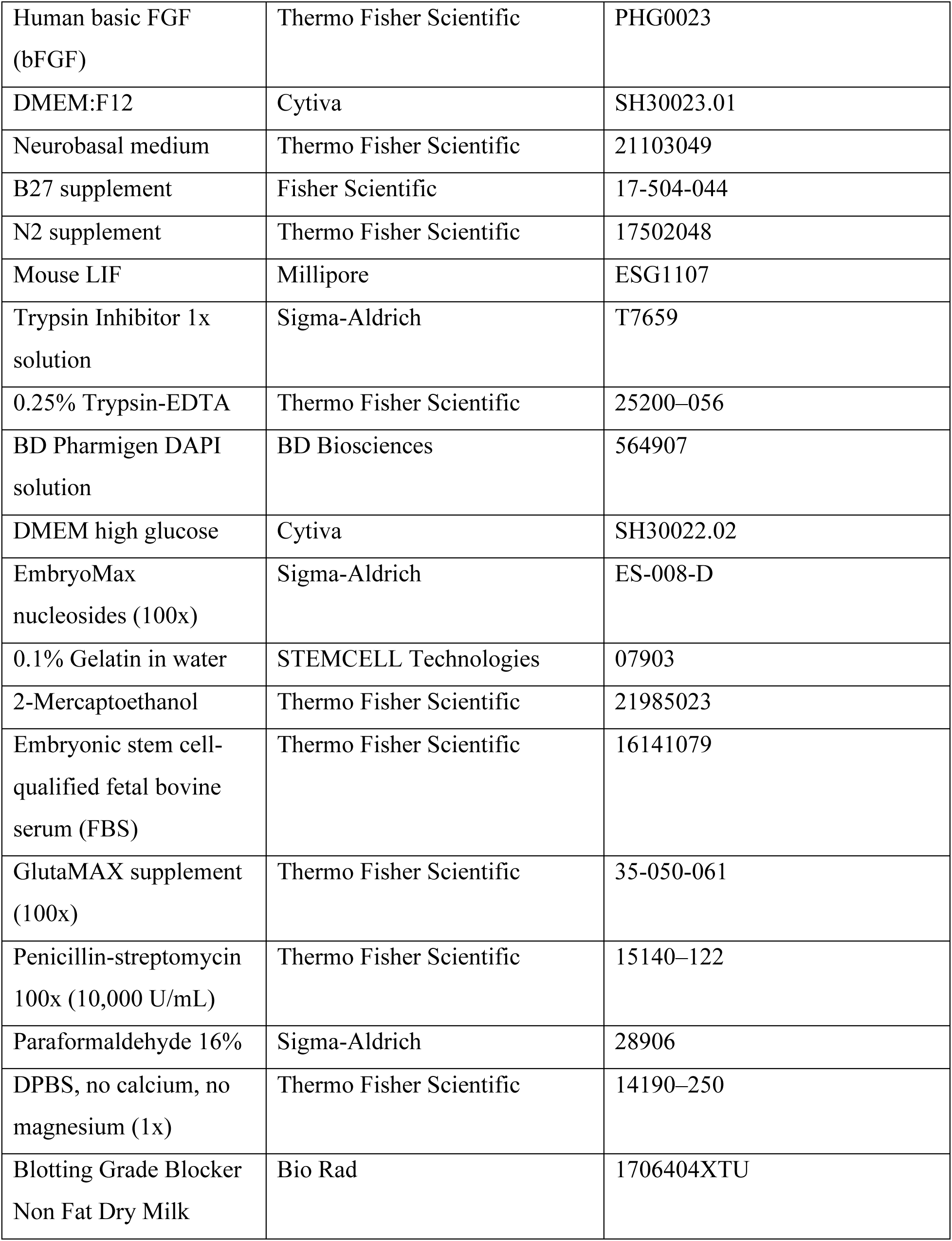
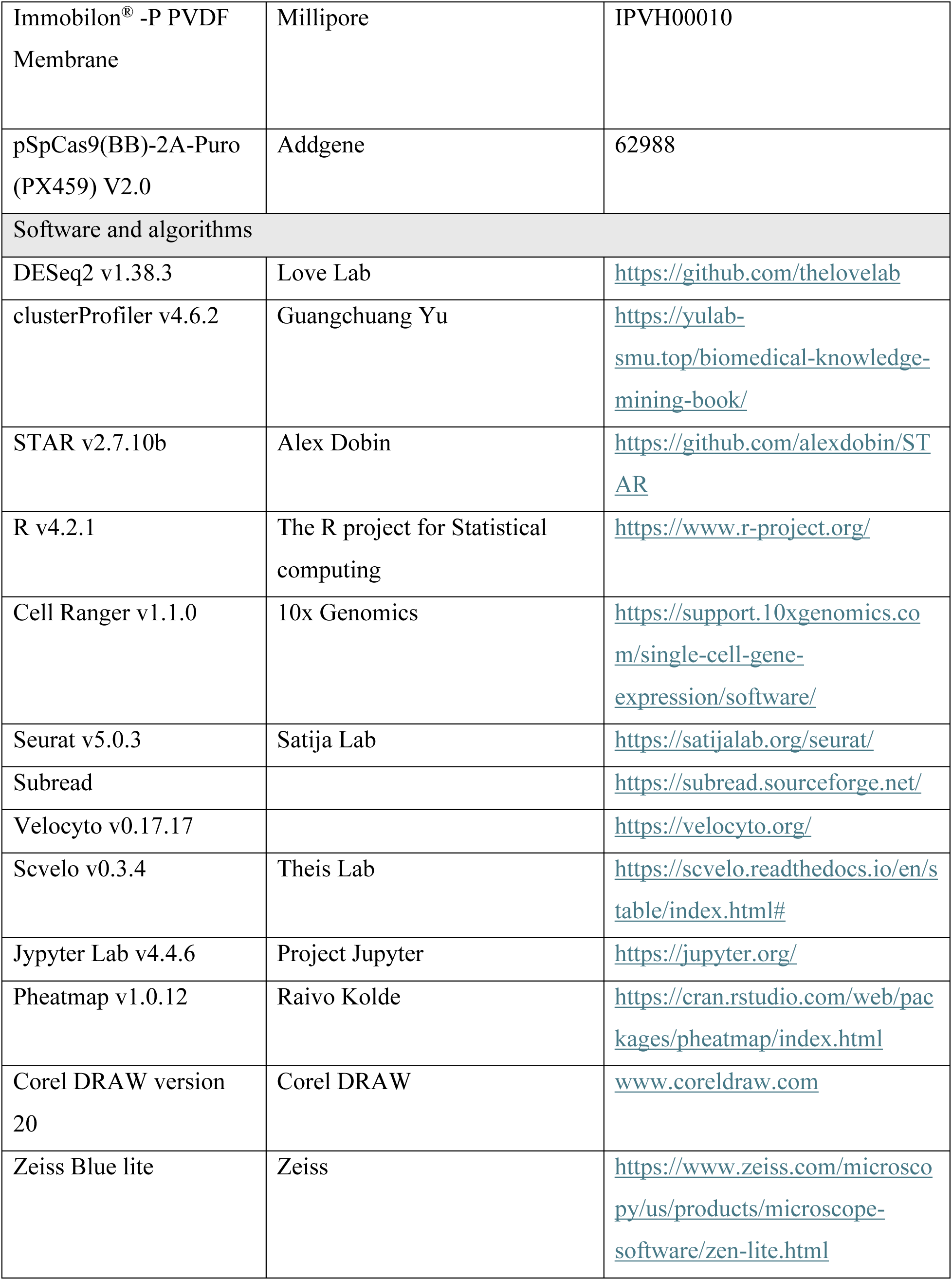
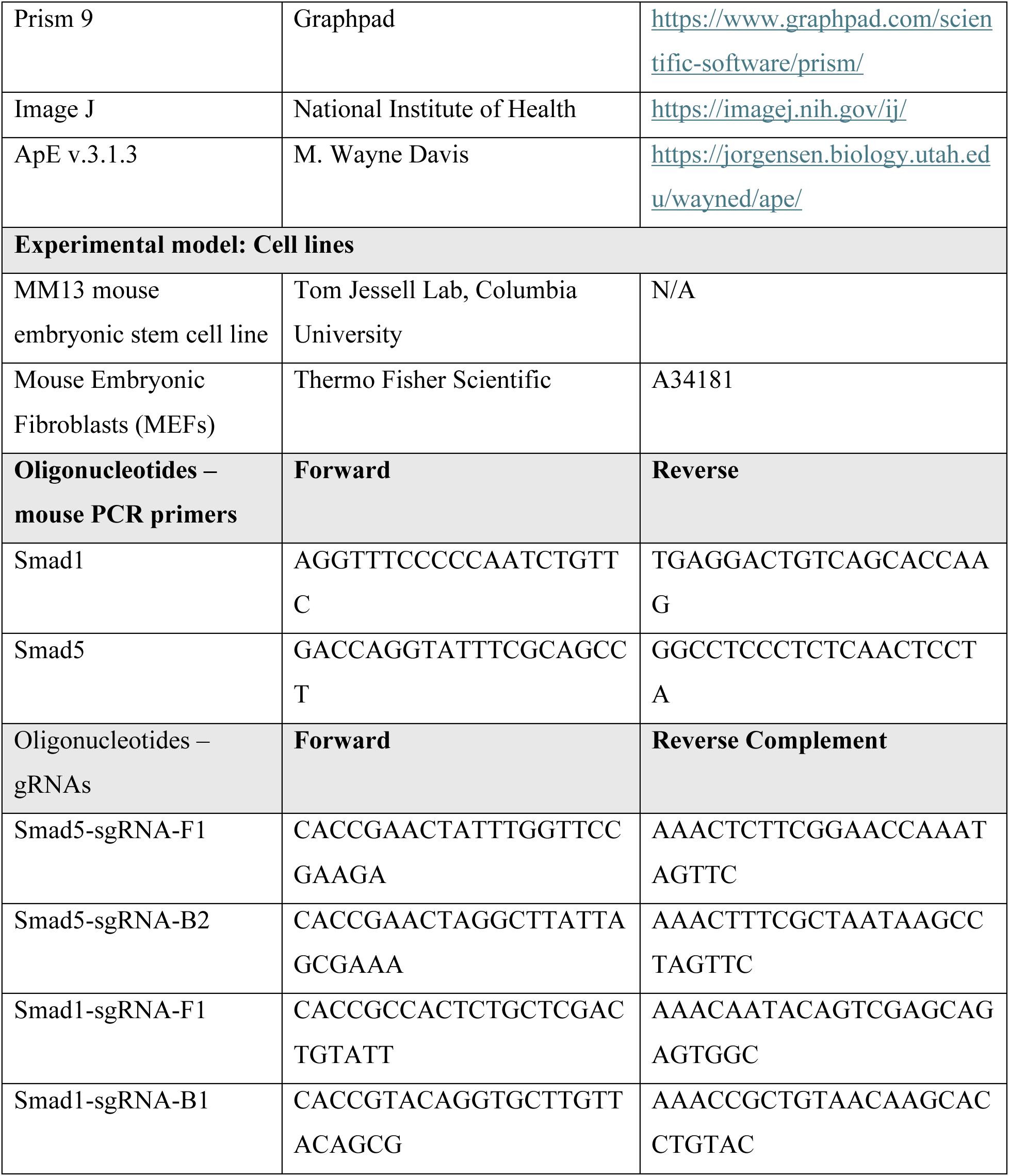

## Results

### Generation of Smad1 and Smad5 knockout mESC lines to investigate dI specification

We have previously established two directed differentiation protocols (RA±BMP4) that derive the complete complement of dIs from mESCs (Fig.1A). These protocols generate dIs that are transcriptionally and functionally indistinguishable from their endogenous counterparts (Gupta et al., 2024; Gupta et al., 2022; Rodriguez et al., 2026), suggesting that together they accurately recapitulate dorsal spinal cord development. Briefly, mESCs are treated with bFGF/CHIR to direct them into a biopotential neuro-mesodermal progenitor (NMP) identity by day 3. The NMPs are then neuralized with retinoic acid (RA) for two days. To generate the dorsal-most dIs, NMPs are additionally treated with BMP4 for 24 hours on day 4. By day 9, post mitotic neural markers are robustly expressed (Myt1l and Tuj1; Fig.1C, 1D, 1F, 1G), together with dP (Pax3; Fig.1C, 1F), and dI markers, which include Lhx2 (dI1), Lhx1/5 (dI2/dI4/dI6), and Lmx1b (dI5) (Fig. 1D, 1G, 1E, 1F). While both protocols generate equivalent numbers of dPs and neurons (Fig. 1I), RA+BMP4 enriches for dI1-dI3, while RA alone results in increased numbers of the intermediate spinal subtypes, dI4-dI6 (Fig. 1C-1J) (Andrews et al., 2017; Gupta et al., 2022).

To begin understanding the role of second messenger signaling in dI specification, we assessed the level and activity of Smad1 and Smad5 in the directed differentiation protocols. Previous studies have shown that Smad8 is not expressed in the developing dorsal spinal cord either *in vivo* (Hazen et al., 2012) or *in vitro* (Gupta et al., 2024). Both Smad1 and Smad5 (Fig. 1K) are present at relatively constant levels along the RA+BMP4 timeline, suggesting BMP addition does not affect Smad protein levels. Rather, the addition of BMP4 at day 4 results in phosphorylated (p; activated) Smad1/5 within 6 hours of treatment (Fig.1K). Elevated pSmad levels persist for 24 hours through day 5. However, by day 7, pSmad levels are comparable across the RA and RA+BMP4 protocols (Fig. 1K, 1L). Together, these data define the temporal window for investigating the transcriptomic changes downstream of canonical BMP signaling that drive dI fate specification and suggest that low level BMP signaling occurs in the RA protocol.

To investigate whether Smad1 and Smad5 have distinct roles mediating BMP-dependent transcriptional programs, we generated Smad1 and Smad5 knockout (Δ) mESC lines. We employed a dual-nicking CRISPR–Cas9 strategy (Ran et al., 2013), designing paired guide RNAs to excise the start codon in exon 2 of the *Smad1* and *Smad5* loci (Fig. 2A). This approach successfully resulted excised exon 2 (Fig. 2B-2D) and thereby generated three independent protein-null mESC lines for each Smad (Fig. 2D). Loss of Smad1 or Smad5 did not alter the mESC morphology during routine passaging, or the expression of key pluripotency markers, such as Oct4, Nanog, and Sox2 (Fig. 2F).

**Figure 2.**
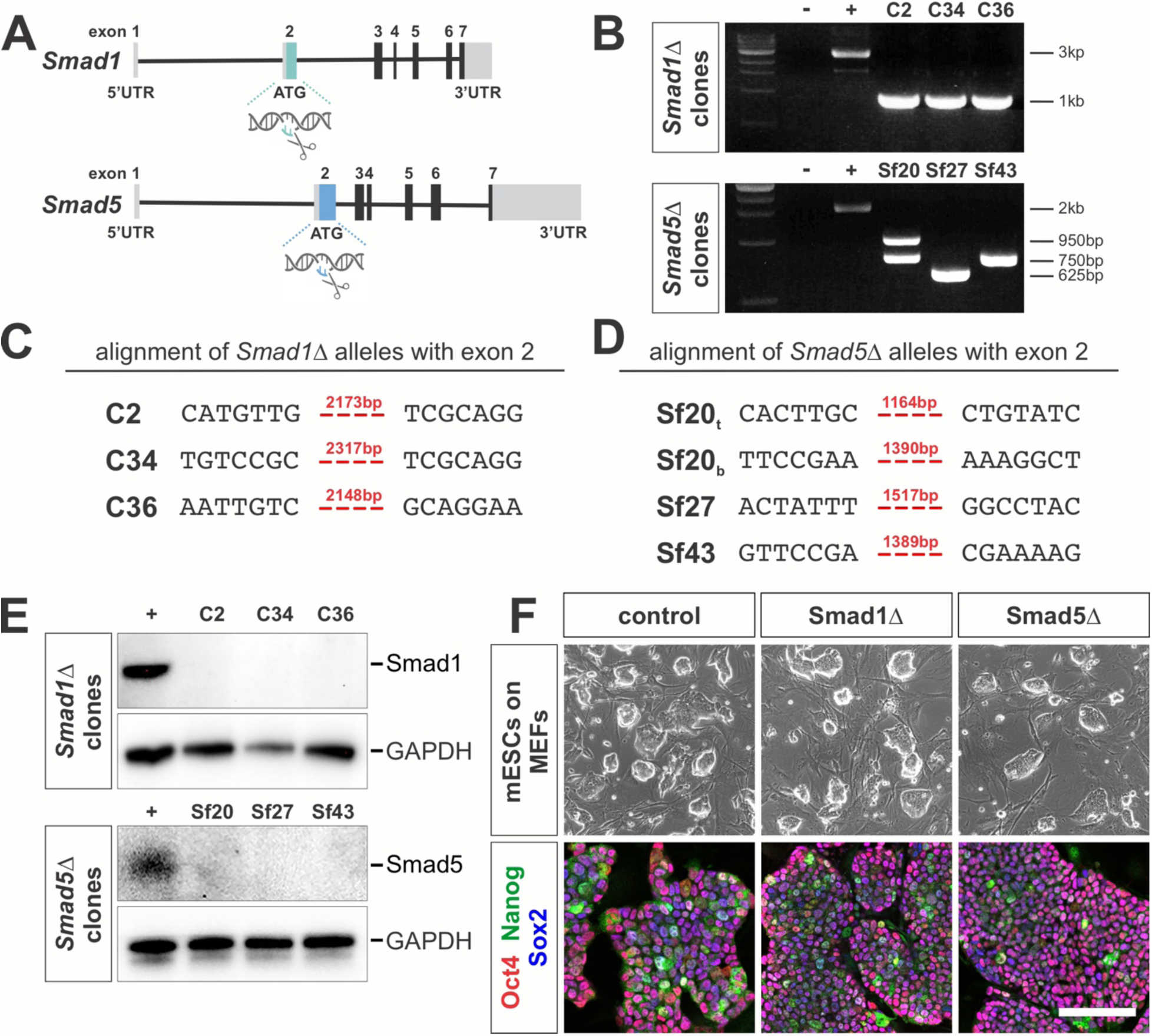
Generation of Smad1 and Smad5 knockout mESC lines (A) CRISPR-Cas9 double nicking gene editing strategy. Guide RNAs flanking exon 2 were used to delete (Δ) the start codon in either *Smad1* or *Smad5* transcript, to thereby disrupt protein translation. (B) Agarose gel electrophoresis of PCR amplicons spanning the targeted region in three independent Smad1Δ lines (C2, C34, C36) and three independent Smad5Δ lines (Sf20, Sf27, Sf43), demonstrated genomic deletions at the xon 2 locus. (C) Exon 2 deletion was confirmed by sequencing and aligning the PCR amplicons to the mouse reference genome, showing that the deletion breakpoints correspond precisely to the gRNA-targeted cut sites. (E) Western analyses validated that all Smad1Δ or Smad5Δ lines contain null alleles. (F) Phase contrast imaging shows that Smad1Δ and Smad5Δ mESCs are morphologically comparable to the isogenic control line (top, F) and continue to express pluripotency markers Oct4, Nanog, and Sox2 (bottom, F). Scale bar: 100µm

### Transcriptomic profiling of WT, Smad1Δ and Smad5Δ lines along the dI differentiation timeline

To identify the gene networks regulated by Smad1 and Smad5 during dI specification, we performed bulk RNA sequencing (Seq) in the Smad1Δ (n=3 independent lines), Smad5Δ (n=3 lines), and isogenic control (n=3 samples) mESC lines at key developmental time points (Fig. 3A) (Gupta et al., 2022). These timepoints were: [1] day 0, pluripotent mESCs; [2] day 3, NMP formation; [3] day 4, after 24 hours of RA treatment; [4] day 4.25 and [5] day 5, 6 and 24 hours respectively after BMP4 treatment, when cells enter a dorsal progenitor (dP) state; and [6] day 9, dI differentiation (n = 75 samples, after library preparation and QC, see Methods for more details).

**Figure 3.**
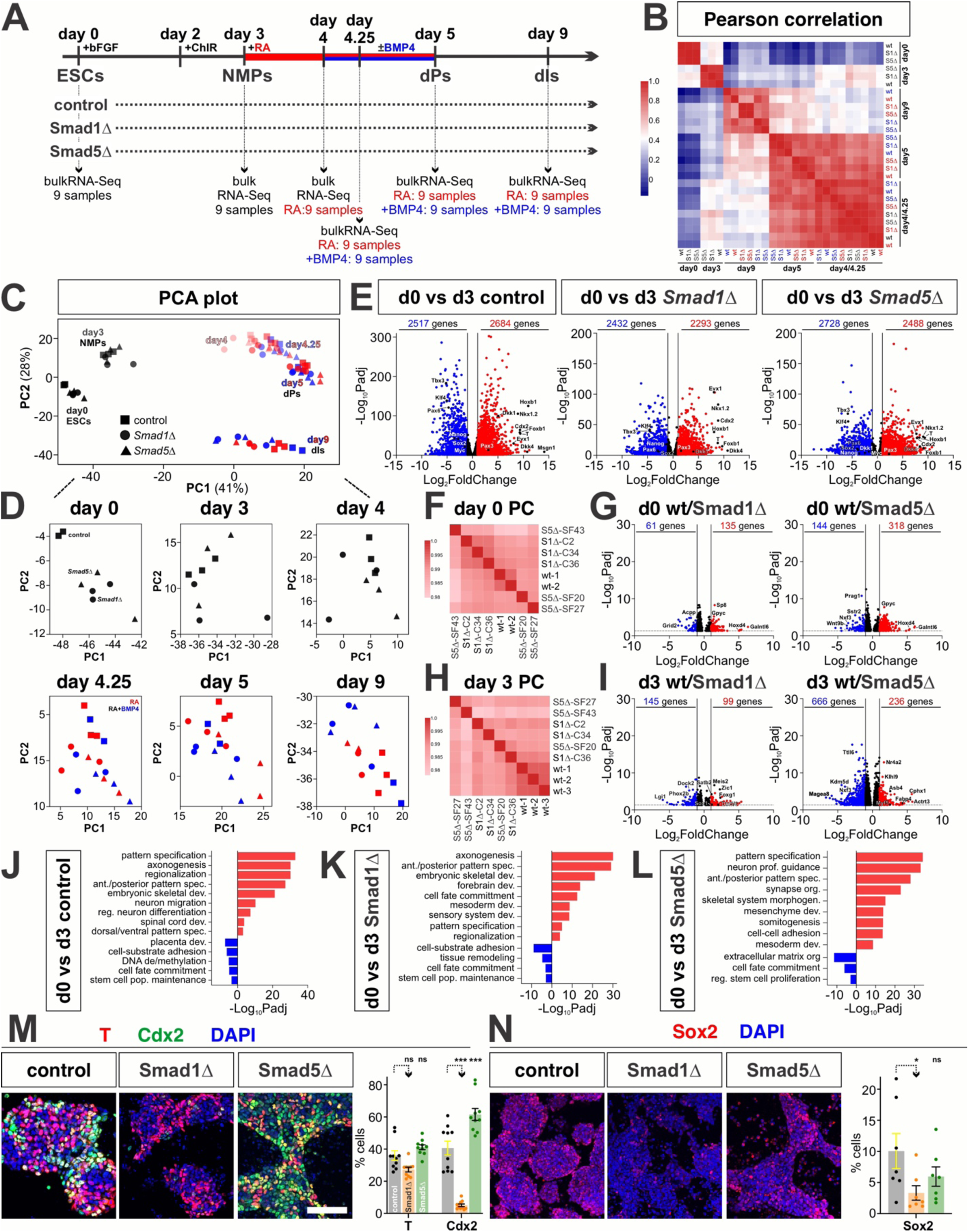
Smad1/5 play opposing roles in NMP fate commitment toward neural and mesodermal lineages at day 3 (A) Timeline for bulk RNA-sequencing experiment (81 samples total). RNA was collected at the indicated developmental time points along the protocol. Three biological replicates were used for each genotype, i.e., from the three independently generated *Smad1Δ* and *Smad5Δ* mESC lines. (B) Pearson correlation of averaged biological replicates at each time point and condition. Samples clustered primarily by developmental stage, with a block of high correlation from 6–24 hours after BMP4 addition, indicating highly similar transcriptional states during this time window. (C, D) Principal component analysis (PCA, C) revealed a shared trajectory as mESCs (day 0) transition towards dorsal interneurons (dIs, day 9). The factor loadings (D) show that Smad1Δ and Smad5Δ samples progressively deviate from the control lines over time, consistent with transcriptional disruptions accumulating as differentiation proceeds. (E) Volcano plots comparing day 0 to day 3 samples show consistent upregulation of NMP-associated genes (*T*, *Cdx2*, *Nkx1.2*) and downregulation of pluripotency genes (*Tbx3*, *Klf4*, *Nanog*) across control, Smad1Δ, and Smad5Δ lines, suggesting all lines successfully acquire NMP fate. (F-I) Pearson correlations (F, G) and volcano plots (H, I) comparing Smad1Δ and Smad5Δ lines to the control at days 0 (F, G) and 3 (H, I). All lines remain highly transcriptionally correlated at both time points; however, differential expression analysis revealed that Smad1Δ cells show increased expression of neural-associated genes (*Zic1*, *Sox1*, *Foxg1*), whereas Smad5Δ cells upregulate mesoderm-associated markers (*Myf5*, *Asb4*, *Fabp4*) relative to controls. (J–L) Gene Ontology (GO) analyses of the significantly up- and down-regulated genes at day 3 (relative to day 0) revealed enrichment for neural and mesodermal biological processes. (M–N) As validation, cultures were immunostained for Brachyury (T, red), Cdx2 (green), Sox2 (red) and DAPI (blue). Decreased numbers of Smad1Δ cells express Cdx2 (M, n=10 images) and Sox2 (N, n=7). In contrast, increased number of Smad5Δ cells express T, suggesting a mesodermal identity. Scale bar: 100µm Significance threshold: log_2_ fold-change –1 > or 1 < and adjusted p-value < 0.05. Probability of similarity between control and experimental groups: * p<0.05, *** p<0.0005; two-way ANOVA

Principal component analysis showed that samples clustered primarily by developmental time point projecting along the expected trajectory from pluripotent to NMP to neural states (Fig. 3C). The Smad1Δ and Smad5Δ lines followed the same overall temporal progression, although their positions were consistently shifted relative to the controls (Fig. 3D), suggesting altered transcriptional states rather than a complete disruption of differentiation. Supporting this conclusion, Pearson correlations showed that Smad1Δ, Smad5Δ, and control samples are highly correlated at day 0 (Fig. 3F), and day 3 (Fig. 3H). The samples segregate most closely according to development time point, rather than genetic background, with differences becoming most pronounced by day 9 (Fig. 3B). Volcano plots comparing day 0 mESCs from the Smad1Δ, Smad5Δ, and control lines also show relatively few differentially regulated genes (Fig. 3G).

### Smad1Δ and Smad5Δ mESCs generate NMPs with altered neuromesodermal potential

We further assessed the consequences of Smad1Δ and Smad5Δ on NMP differentiation by comparing the day 3 transcriptome from each line to control day 0 mESCs. Control, Smad1Δ and Smad5Δ cells all downregulated the pluripotency genes, *Tbx3*, *Klf4*, and *Nanog*, and upregulated NMP markers, including *Nkx1.2*, *Cdx2*, *T,* and *Evx1* (Fig. 3E). Gene Ontology (GO) analysis of significantly upregulated genes at day 3 showed enrichment for terms associated with both mesodermal and neurogenic programs (Fig. 3J-3L). Together, these data indicate that the loss of Smad1 or Smad5 does not prevent the formation of NMPs.

While Smad1Δ, Smad5Δ, and control samples were highly transcriptionally correlated at day 3 (Fig. 3H), some transcriptional differences relative to control were observed (Fig. 3I). Smad1Δ cells showed increased expression of *Zic1*, *Foxg1*, and *Sox1*, genes involved in early neural development and neural progenitor maintenance. In contrast, Smad5Δ cells showed increased expression of mesoderm-associated genes, including *Myf5*, *Asb4*, and *Fabp4* (Fig. 3I). Thus, although the Smad mutants attain an NMP identity, they may be biased towards distinct lineage trajectories. Supporting this hypothesis, immunohistochemistry showed that more Smad5Δ NMPs express Cdx2 protein than the control, consistent with enhanced mesodermal priming (Fig. 3M). In contrast, Smad1Δ NMPs show decreased Cdx2 and Sox2 levels, suggesting they are primed away from a mesodermal trajectory (Fig. 3N).

### Stimulating NMPs with BMP4 during dI differentiation increases chromatin accessibility

We next sought to assess the chromatin landscape from day 4-5 to better understand the developmental events that occur after recombinant BMP4 is added to the differentiation media. Given that chromatin accessibility changes precede transcriptional outputs, this analysis also provided a framework for interpreting subsequent gene expression differences. Assay for Transposase Accessible Chromatin (ATAC)-Seq was performed at days 4, 4.25, and 5 capturing chromatin accessibility changes 6 and 24 hours after adding BMP4 (Fig. 4A). Combining peaks across all conditions and time points generated a unified ATAC-Seq peak set of 141,592 peaks. ∼46% of peaks mapped to intergenic regions, indicating that the largest fraction of accessible chromatin corresponds to distal regulatory elements (Fig. 4B).

**Figure 4.**
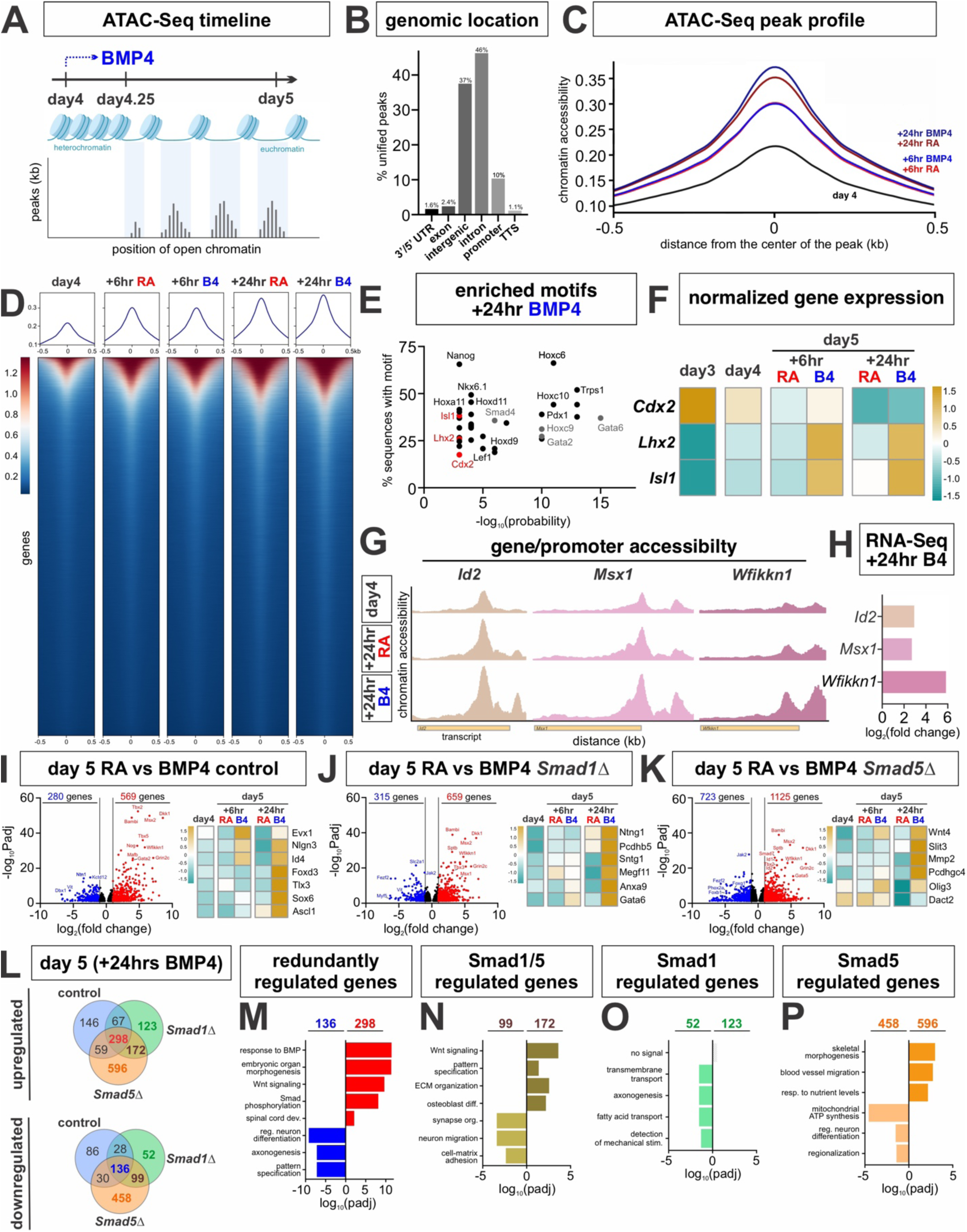
ATAC-Seq analyses from days 4- 5 reveal divergent epigenomic and transcriptional roles for Smad1 and Smad5 downstream of BMP4 treatment (A) Schematic timeline of ATAC-Seq studies to assess the chromatin accessibility changes before (day 4) and after (day 4.25 and 5) BMP4 treatment. Three biological replicates were used for each time point (control line only; n=9 samples). (B) ATAC-Seq peaks across samples were merged to form a unified peak set (n = 141,592). The majority of peaks fall in intronic (46%) and intergenic (37%) regions and together account for ∼83% of all peaks. (C, D) Accessibility profiles of differentially accessible regions showed progressive increases in chromatin accessibility from day 4 to day 5. While there were no significant differences between BMP4-treated and control samples at 6 hours, clear accessibility changes emerge by 24 hours. (E) Motif enrichment analysis identified 37 transcription factor motifs significantly enriched in regions with increased chromatin accessibility following 24 hours of BMP4 treatment. (F) Heatmap examining the expression of key transcription factors identified in (E), shows BMP4 treatment rapidly upregulates *Lhx2* (dI1), and *Isl1* (dI3) expression. However, *Cdx2* (mesoderm) is progressively downregulated, even though Cdx2 motifs remain accessible 24 hours after BMP4 treatment. Thus, BMP4 treatment may selectively stabilize pre-existing accessible regions. (G, H) Similarly, by 24 hours after BMP4 addition, chromatin accessibility is increased at the promotors of established BMP-regulated genes, such as *Id2* and *Msx1*, with concomitant transcriptional upregulation. This analysis also identified novel regulators of dorsal fate specification, such as *Wfikkn1,* a secreted ligand that can sequesters BMP4, and be part of a negative feedback mechanism that limits BMP signaling. (I–K) Building on the localized chromatin remodeling observed 24 hours after BMP4 treatment, differential gene expression was examined in the control, Smad1Δ, and Smad5Δ bulk RNA-Seqdatasets at day 5. Volcano plots show that the most upregulated genes across the lines are shared canonical BMP target genes. Heatmaps further identify genes uniquely regulated in each line, revealing both Smad-specific and shared transcriptional responses to BMP4. (L) Venn diagram showing the overlap of genes up or downregulated after 24 hours of BMP4 treatment between Smad1Δ, Smad5Δ and control cells. Loss of Smad5 consistently affects a larger subset of genes. (M-P) GO analyses of subsets of genes taken from the Venn diagram (L). The 298 genes redundantly upregulated by Smad1 and Smad5 show enrichment for BMP signaling pathway terms (M). Shared transcriptional changes in Smad1Δ and Smad5Δ cells (172 upregulated, 99 downregulated genes) are associated with increased extracellular matrix organization and decreased neuron migration and cell adhesion (N). Strikingly, there are relatively few transcriptional changes unique to the loss of Smad1 (O), whereas Smad5Δ leads to more extensive transcriptional remodeling associated with non-canonical BMP functions, such as altered regulation of bioenergetics (P).

To assess whether BMP4 alters chromatin accessibility, we examined the ATAC-Seq peak profile across developmental time and treatment conditions. At 6 hours, the chromatin accessibility profiles were essentially the same between RA- and RA+BMP4-treated samples, indicating minimal early remodeling. However, by day 5, the RA+BMP4-treated samples exhibited increased accessibility relative to the RA-control (Fig. 4C). These changes were restricted to discrete regulatory elements, indicating that BMP4 induces localized rather than global chromatin remodeling (Fig. 4D).

### Chromatin accessibility profiling identifies gene targets of BMP4 signaling

ATAC-Seq peaks showing increased accessibility after 24 hours of BMP4 treatment compared to RA-treated controls (log2FC >0.5) were defined as BMP4-responsive regions. Of the 154 BMP4 responsive regions, ∼36% contained the Smad Binding Element (SBE) “GTCT” core sequence (Shi et al., 1998). HOMER motif enrichment analysis identified 53 transcription factor binding motifs associated with BMP4-induced chromatin accessibility changes, 37 of which are present in >15% of target sequences, and include *Gata2*, *Gata6*, *Isl1*, *Lhx2* and many *Hox* genes (Fig. 4E), thereby identifying new and established transcription factors in the response to BMP4. *Lhx2* and *Isl1* expression marks dI1/dI3 identities respectively (Alaynick et al., 2011) and are also upregulated after 6 hours of BMP4 treatment (Fig. 4F). These factors are an early response to BMP4 and may thus drive the initial regulatory programs underlying BMP-dependent chromatin remodeling and gene activation in the dI1/3 subtypes. 32 transcription factor motifs enriched in BMP4-responsive chromatin were not concurrently transcriptionally upregulated, suggesting they reflect a residual chromatin footprint of prior transcription factor activity. For example, *Cdx2* expression is highly expressed in day 3 NMPs, but downregulated by day 5 (Figs. 3M, 4F); yet its motif remains enriched 24 hours after BMP4 treatment, suggesting that BMP4 can selectively stabilized pre-existing accessible regions.

We next sought to identify genes showing concordant promoter accessibility and transcriptional upregulation after BMP4 treatment. 16 promoter regions with significantly increased accessibility in 24-hour BMP4-treated samples were identified, mapped to their closest genes, and validated against bulk RNA-Seq data from days 4 and 5 (Fig. 4G, 4H). This analysis identified *Id2*, *Msx1*, and *Wfikkn1* which all display increased chromatin accessibility proximal to the transcription start site, consistent with promoter remodeling (Fig. 4G). Notably, *Id2* and *Msx1* are established transcriptional targets of BMP signaling (Hollnagel et al., 1999; Le Dreau et al., 2018; Liem et al., 1995) supporting the model that BMP4 promotes their activation through localized, promoter-associated chromatin accessibility changes. Wfikkn1 (WAP, follistatin/kazal, immunoglobulin, kunitz, and NTR domain–containing) has not previously implicated in dorsal spinal cord development; however, it encodes a secreted multidomain protein that can bind BMP4 and antagonize GDF8 and GDF11 (Kondas et al., 2008; Szlama et al., 2010). Thus, BMP4 may dynamically induce *Wfikkn1* expression to establish a negative feedback mechanism that fine-tunes BMP ligand availability during neural differentiation.

### Smad1 and Smad5 have shared and distinct roles downstream of BMP4 signaling

ATAC-Seq analysis revealed minimal chromatin remodeling at 6 hours (day 4.25) but robust, localized regulatory changes at 24 hours following BMP4 treatment (Fig. 4C, 4D). We therefore focused on this later timepoint to assess transcriptional differences in control, Smad1Δ and Smad5Δ lines in the day 5 bulk RNA-Seq data, 24 hours after BMP4 addition. For each genotype, BMP4 treated samples were compared to matched RA-only controls to identify BMP-dependent transcriptional responses, and the overlap between up- or downregulated genes was determined for the Smad1Δ, Smad5Δ and control cells (Fig. 4L).

298 genes were upregulated in all three lines, accounting for 45% and 26% of the genes upregulated in Smad1Δ and Smad5Δ samples (Fig. 4L). This shared gene set included established BMP targets, such as *Id1*, *Id2*, and *Wfikkn1* (Fig. 4I, 4L), and GO analysis returned terms including “response to BMP” and “Wnt signaling pathway,” consistent with our previous findings (Gupta et al., 2022) (Fig. 4M). Since either Smad1 or Smad5 are required for this response, the R-Smads appear to act redundantly to mediate these BMP-responsive transcriptional programs. In contrast, 67 and 59 genes were specifically upregulated in control/Smad1Δ cells and control/Smad5Δ cells respectively (Fig. 4L), identifying candidate targets selectively regulated by Smad1 or Smad5 (heatmaps, Fig. 4J and Fig. 4K).

We further identified 172 upregulated and 99 downregulated genes expressed in Smad1Δ and Smad5Δ, but not control, BMP4-treated cells (Fig. 4L). The presence of these genes exclusively in the Smad1Δ and Smad5Δ datasets suggests that loss of either R-Smad disrupts shared transcriptional mechanisms. A GO analysis of these genes implicated pathways involved in extracellular matrix organization, synapse organization, and neural migration (Fig. 4N). Finally, there were relatively few (175) uniquely regulated genes in the Smad1Δ cells, which yielded minimal GO term enrichment (Fig. 4O). In contrast, there were substantially more (1054) uniquely regulated genes in Smad5Δ cells (Fig. 4L). GO analysis suggests these Smad5-dependent genes regulate non-canonical or context-dependent functions (Fig. 4P), including bioenergetic homeostasis as previously shown (Fang et al., 2017). Taken together, these findings support a model in which Smad1 and Smad5 play shared and distinct transcriptional roles mediating the transition from the NMP to dP state, with Smad5 having a broader role regulating the response to BMP4.

### Smad1 and Smad5 have distinct roles in dorsal spinal lineage specification

To assess whether Smad1 and Smad5 are differentially required for dorsal lineage specification, we generated single-cell (sc) RNA-Seq atlases for Smad1Δ, Smad5Δ, and control cells at day 9 of the RA±BMP4 differentiation protocol. To enable accurate comparison across the six datasets [RA-control, n=13,822; RA-Smad1Δ, n=11,157; RA-Smad5Δ, n=10,616; RA+BMP4-control, n=8,276; RA+BMP4-Smad1Δ, n=10,875; and RA+BMP4-Smad5Δ, n=7,944], they were integrated into a single combined transcriptional atlas (Fig. 5A). Given the Smad-dependent alterations in NMP specification (Fig. 3), we first assessed the proportion of *Foxc1^+^*, *Twist1*^+^, *Meox2*^+^ mesodermal and *Pax3*^+^, *Myt1l*^+^, *Msx1*^+^ neural derivatives generated in the three lines (Fig. 5B, 5C, Figure S1). In the RA condition, ∼90% of cells were classified as neural across genotypes (Fig. 5D). BMP4 addition resulted in ∼25% of cells following a non-neural (mesodermal) trajectory in both the control and Smad1Δ samples (Fig. 5E). This proportion was further expanded in the Smad5Δ group, which had ∼40% non-neuronal population (Fig. 5E), including both mesoderm, and a marked expansion of *Sox17^+^*, *Col4a2^+^*, *Gata4^+^*, *Foxa2^+^* extraembryonic endoderm-like cells. These findings support a model in which Smad5 biases NMPs towards neural fates, at the expense of non-neural fates, including mesoderm induction (Fig. 5J).

**Figure 5.**
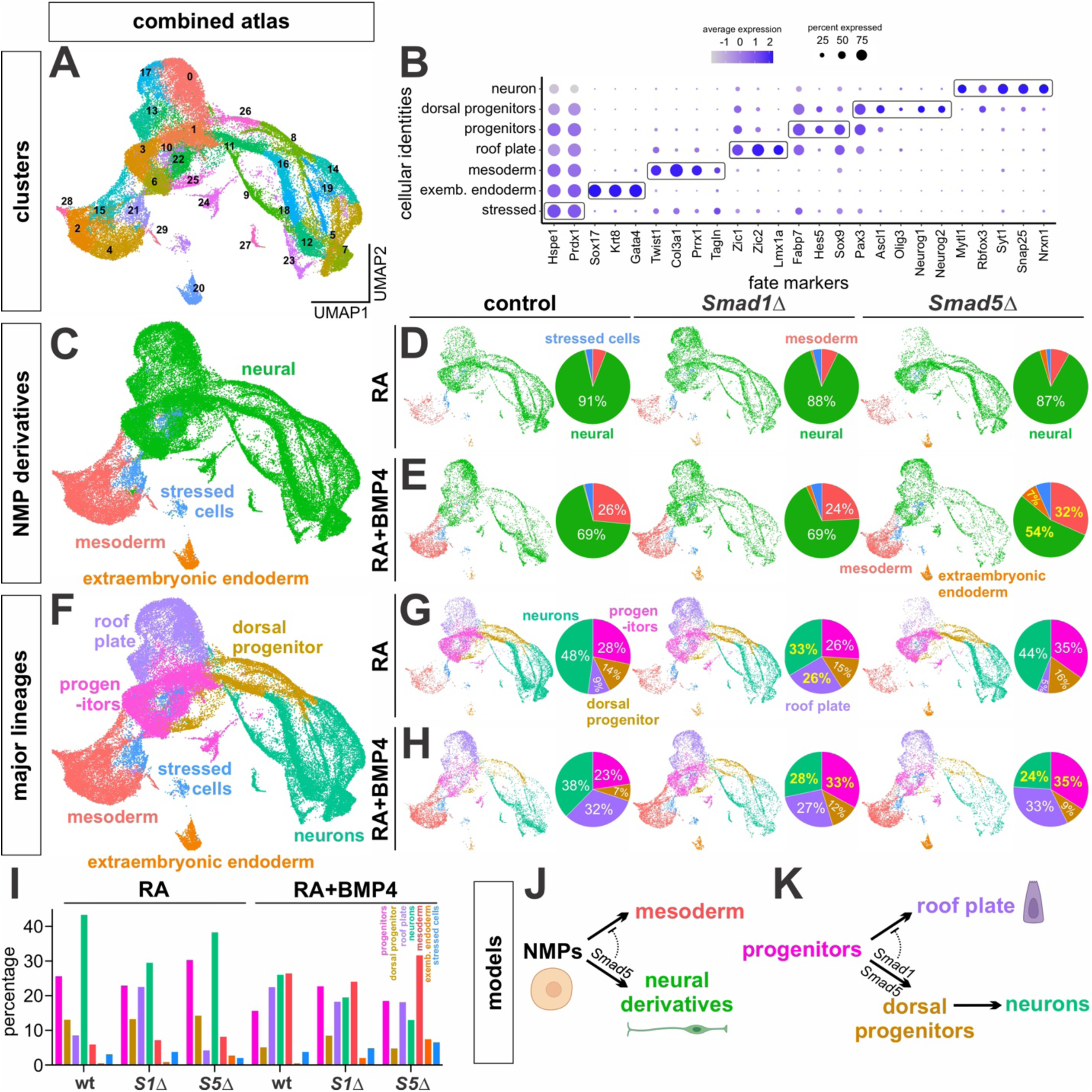
The loss of Smad5 alters lineage specification during dI *in vitro* differentiation (A) Integrated UMAP atlas of scRNA-Seq datasets from RA±BMP4 directed differentiations of the Smad1 (C36 line), Smad5 (SF43 line) and isogenic control lines at day 9 (n=6 datasets). Clustering of the data yields 30 transcriptionally distinct cell clusters. (B) Dot plot showing the expression levels and identity of marker genes used to define the different lineage identities derived from NMPs. (C) Integrated UMAP atlas depicting the mesodermal (coral) neural (green), and extraembryonic endoderm (orange) lineage trajectories. Stressed cells are labeled in blue. (D, E) Pie charts and accompanying sample-specific UMAP atlases quantifying the proportion of neural, mesodermal, and extraembryonic endoderm cell derivatives across sample groups. There are no major cell fate specification differences in the RA protocol, however in the RA+BMP4 protocol, the mesodermal and extraembryonic endoderm populations are expanded specifically in the Smad5Δ line. (F) Integrated UMAP atlas identifying the six major lineage groups, and the stressed cell population. (G, H) Pie charts and accompanying sample-specific UMAP atlases quantifying the proportions of progenitor, roof plate, dorsal progenitor, and neural cell populations. In the RA protocol, the roof plate population is expanded at the expense of the neural population in the Smad1Δ line. In the RA+BMP4 protocol, both the Smad1Δ and Smad5Δ lines have the same phenotype: the progenitor population is increased, and the number of neurons concomitantly decreased. (I) Bar chart summarizing the percentage of the major cell lineages differentiated from control, Smad1Δ and Smad5Δ line using the RA ± BMP4 protocols. (J, K) Models summarizing the roles of Smad1 and Smad5 directing the specification of NMP derivatives. Smad5 plays the earliest role in lineage specification, preventing NMPs from entering a mesodermal trajectory (J). As differentiation proceeds, both Smad1 and 5 are required for progenitors to progress towards a neural fate (K); Smad1 additionally prevents progenitor cells from being directed towards a roof plate fate.

We then subdivided the neural category into broad lineage divisions – cycling progenitors, roof plate, dorsal progenitors, neurons - based on the expression of canonical marker genes (Fig. 5B, 5F). In the RA+BMP4 condition, where R-Smads are activated in response to extrinsic BMP4 (Fig 1N), progenitors accumulate at the expense of neurons in both R-SmadΔ lines, compared to control (Fig. 5H). However, in the RA condition, where R-Smads are activated by intrinsic BMP signaling (Fig 1L), control and Smad5Δ lines show similar cellular proportions, while roof plate cells accumulate at the expense of neurons in the Smad1Δ line (Fig. 5G). Together, these data suggest that both R-Smads are required for progenitors to initiate the differentiation process, but then Smad1 specifically suppresses them from progressing to a roof plate fate (Fig. 5K).

Finally, we assessed the requirement for Smad1 and Smad5 in the specification of dPs and dIs. The scRNA-Seq data for dPs and neurons were subsetted and reclustered to generate an integrated neural atlas for Smad1Δ, Smad5Δ, and control cells (Fig. 6A). Annotation using established dP and dI marker genes (Fig. 6B) confirmed that the RA±BMP4 protocols direct the complete complement of dP (dP1-dP6), and dI (dI1- dI6) populations (Fig. 6C, 6F). dP phenotypes were specifically observed in the RA protocol, where the R-Smads have opposing effects. Smad5Δ results in the loss of dP1-dP3, whereas Smad1Δ increases the proportion of dP1 and dP2 (Fig. 6D, 6I), suggesting the R-Smads function antagonistically in dP specification. dI phenotypes were observed in both protocols (Fig. 6J). Most notably, the dorsal-most dIs, especially the dI1s, were markedly and consistently lost in the Smad5Δ condition relative to control, with a corresponding increase in the intermediate dI fates (Fig. 6H, 6I). dI1s were also lost in the Smad1Δ condition, but only in the RA+BMP4 protocol, perhaps reflecting a requirement for Smad1 when BMP4 signaling is amplified. The dI1 phenotype was validated by immunohistochemistry, confirming a significant reduction in Lhx2^+^ dI1 cells specifically in the Smad5Δ line (Fig. 6K, 6L). Together, these data support a model where the R-Smads play distinct roles in dI fate specification. Smad5 plays the dominant role, acting reiteratively to specify the dorsal-most dI fates (Fig. 7M).

**Figure 6.**
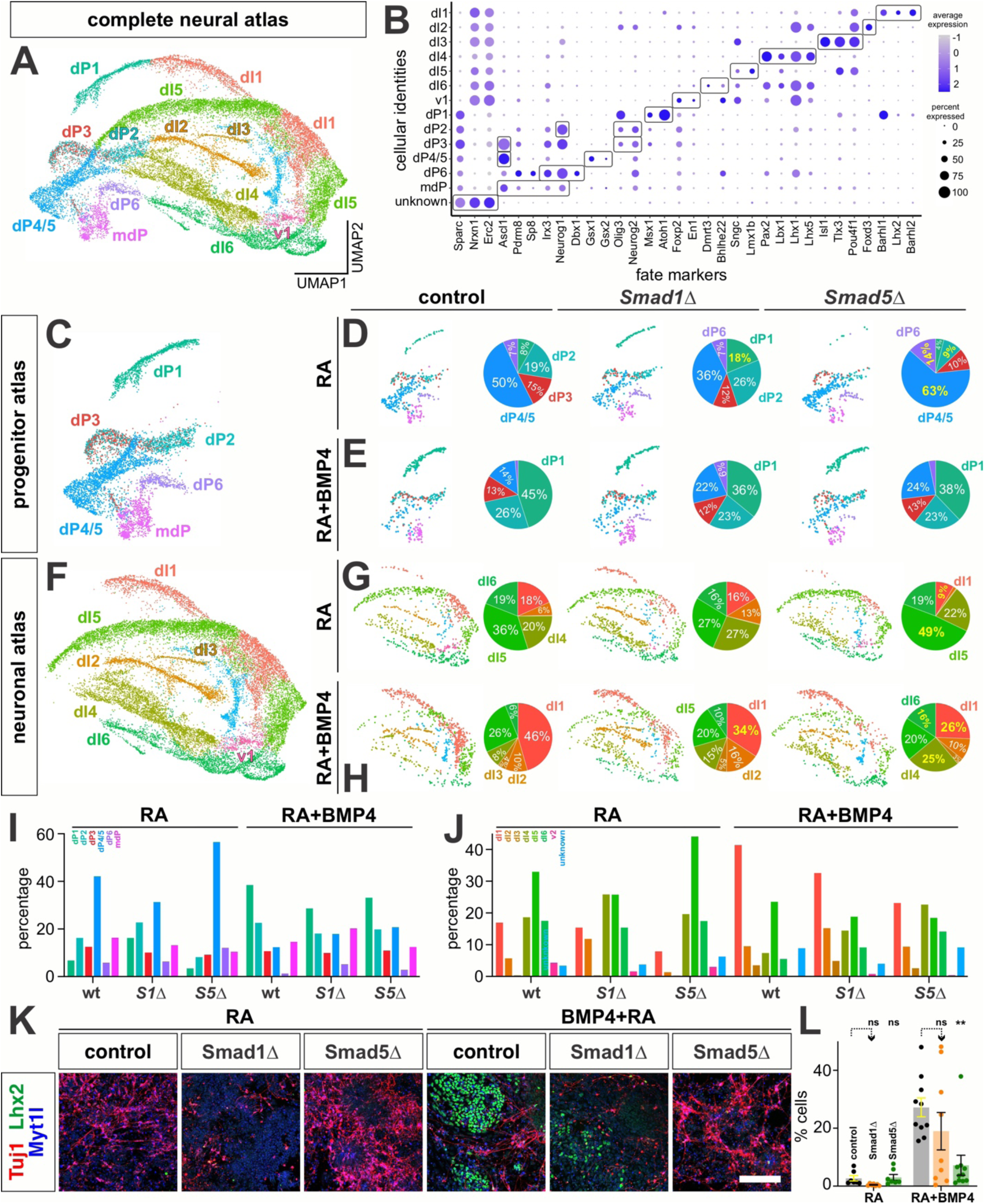
Smad1 and Smad5 are differentially required to specify dI cell fates (A) Cells annotated as dorsal progenitors and neurons were re-clustered to generate an integrated neural UMAP atlas, capturing the dP1–dP6 and dI1–dI6 trajectories. Based on RNA velocity analyses (Figure 7), a population located at the periphery of the atlas, that appears to give rise to dP2–dP6, was annotated as "mdP" (multipotent dorsal progenitor). (B) Dot plots showing the expression levels and identities of marker genes used to assign neural identities (C-H) UMAPs of the integrated dP (C) and dI (F) atlases, and sample-specific UMAP atlases for the control, Smad1Δ and Smad5Δ lines in the RA ± BMP4 protocols (D, E, G, H). The RA protocol robustly generates dP/dIs from the intermediate spinal cord (dI4-dI6), with more modest numbers of dorsal-most dP/dIs, i.e. dI1-dI3. Under these conditions, the Smad5Δ line shows impaired formation of dP1s and dP2s (D), with correspondingly reduced numbers of dI1s and dI2s (G). In contrast, the Smad1Δ line shows increased numbers of dP1/dP2s (D), but no corresponding change in the number of dI1/dI2s (G). In the RA+BMP4 protocol, both the Smad1Δ and Smad5Δ lines show decreased numbers of dP1/dP2s (E) and dI1/dI2s (H). (I, J) Bar charts summarizing the percentage of dP (I) and dI (J) found in control, Smad1Δ and Smad5Δ cell populations using the RA ± BMP4 protocols. (K, L) As validation, cultures were immunostained for Tuj1 (axons, red), Lhx2 (dI1, green), and Myt1l (neurons, blue). As previously observed (Andrews et al., 2017; Gupta et al., 2022), significant numbers of dI1s are only generated in the RA+BMP4 protocol (L). Supporting the transcriptional analysis, Lhx2^+^ dI1s are most reduced in the Smad5Δ differentiation (n=10 images), compared to either the Smad1Δ (n=9) or control (n=10) differentiation. Scale bar: 100µm Probability of similarity between control and experimental groups: ** p<0.005; two-way ANOVA

**Figure 7.**
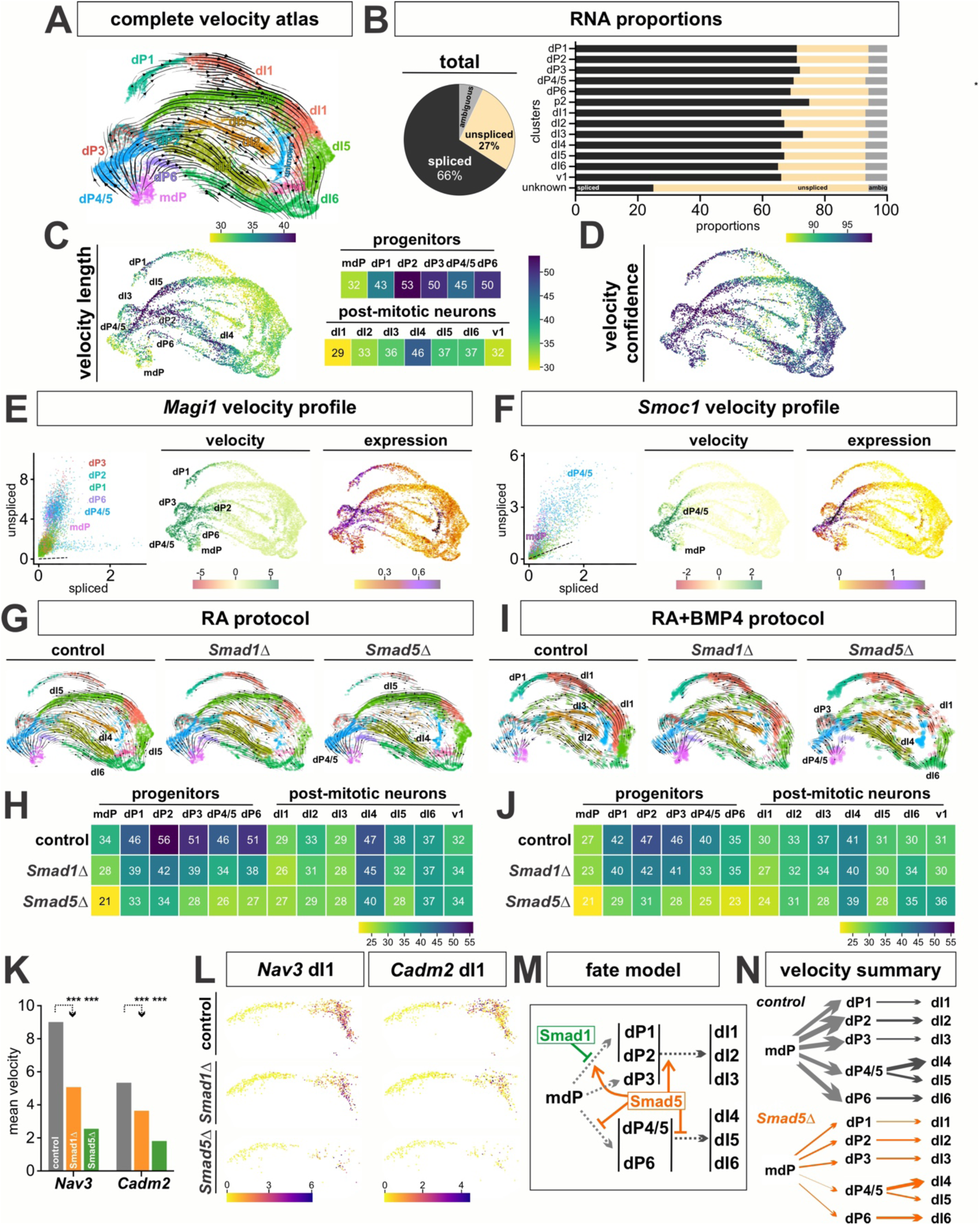
Smad5 is required for normal transcriptional dynamics across all dI lineages (A) RNA velocity analysis of the integrated neural atlas was performed to infer the direction and dynamics of dP cell state transitions. The streamlined velocity vector field shows that each dP/dI subtype follows a distinct developmental trajectory. The dI2-dI6 subtypes stem from a common mdP population, which bifurcates into specific dP subtypes that resolve into their respective dI populations. (B) RNA velocity metrics showing the proportion of spliced vs unspliced transcripts in the complete atlas as well as the individual clusters. (C, D) Velocity magnitude (C) and velocity confidence (D) values from the control RA±BMP4 dataset projected onto the UMAP embedding show generally increased RNA velocity in the dP subtypes relative to the dI populations. A notable exception is the dI4 subtype, which maintains high RNA velocity along its entire trajectory. (E-F) Phase portraits of two genes exhibiting differential RNA velocity in the control RA±BMP4 dataset. *Magi1* expression shows positive velocity across all dP populations (E), while *Smoc1* is specifically positive in the dP4/dP5 population. (G-J) Streamlined representation of the velocity vector fields in control, Smad1Δ and Smad5Δ UMAP atlases in RA±BMP4 protocols (G, I). Heatmaps of the average velocity-length values demonstrates that Smad5Δ results in dramatically reduced velocity in all dP subtypes, in both RA±BMP4 protocols (H, J). (K) Velocity magnitude per gene was quantified for the dI1 population across the control, Smad1Δ and Smad5Δ lines. *Nav3* and *Cadm2* exhibited the highest reductions in velocity, which was most pronounced for the Smad5Δ line. (L) *Nav3* and *Cadm2* expression is progressively diminished in the Smad1Δ and Smad5Δ dI1s, suggesting the reduced velocity magnitude reflects impaired progression of Nav3^+^/Cadm2^+^ dI1s. (M) Smad5 is required reiteratively across the specification of dI trajectories. It is required for the specification of the dorsal-most progenitor (dP1/dP2) and neuronal (dI1-dI3) populations, at the expense of the intermediate dI populations (dI4–dI6). Smad1 acts more narrowly, limiting the pool of dP1s. (N) RNA velocity analysis shows that dP clusters have broadly increased velocity compared to dIs, with the exception of dI4 lineage. We propose Smad5 is the principal mediator of the transcriptional programs that maintain progenitor activity and regulate their progression into differentiated dI states.

### RNA velocity analysis identifies genes governing dP state transitions

To further dissect the role of the R-Smads, we performed an RNA velocity analysis on our integrated neural atlas to infer the direction and dynamics of dI state transitions. The resulting global velocity field showed that dPs directionally flowed into their respective dI lineages in the integrated atlas, reflecting the expected developmental progression (Fig. 7A). Unexpectedly, a transcriptionally distinct progenitor cluster (Fig. 6B) was observed to give rise to dP2-dP6, suggesting this represents a pool of multipotent dorsal progenitor (mdP) cells (Fig. 7A). We next examined the velocity magnitude for the control cells (Fig. 7C), to assess how quickly the transcriptome is changing across different subtypes during dI differentiation. The dP populations exhibited the highest velocity lengths, consistent with their being in a transitioning state (Fig. 7C). By contrast, the dI populations have reduced transcriptional dynamics, consistent with a differentiated state, with exception of the dI4 and early dI5 trajectories, which retain elevated velocity lengths (Fig. 7C). Thus, dI4/dI5s may remain in a dynamic state, as they activate subtype-specific transcriptional programs.

To identify genes that drive or mark these cell state transitions, we performed velocity-based gene ranking across the control clusters. This analysis revealed cluster-specific genes where their transcriptional dynamics are strongly correlated with velocity directionality. *Magi1* and *Smoc1* were identified as two of the top-ranked velocity genes. *Magi1* is broadly expressed across multiple dP clusters (Fig. 7E), and may have a general role regulating cell polarity and cell-cell adhesion (Stetak and Hajnal, 2011). Conversely, *Smoc1* expression is specific to the dP4/5 population, suggesting more specific role in this lineage (Fig. 7F). *Smoc1* has previously been identified a dP5 marker gene in an independent pseudotime analysis (Gupta et al., 2024), and acts as a BMP agonist, although this activity depends on context (DeGroot et al., 2023; Thomas et al., 2017). Smoc1 may thus act through BMP signaling to maintain velocity magnitude in transitioning dP5s. Together, this RNA velocity analysis captures biologically relevant transcriptional dynamics underlying dI state transitions.

### Smad5 controls transcriptional dynamics across dP and dI clusters

Having established the baseline velocity dynamics within a control atlas, we investigated whether these transcriptional programs are altered in Smad1Δ or Smad5Δ cells. We quantified mean velocity length for all dorsal populations in the RA±BMP4 protocols and found that loss of the R-Smads results in reduced velocity magnitudes across the dP clusters, with the most marked decreases in Smad5Δ cells (Fig. 7G-7J). Notably, the dP4-dP6 populations, previously thought to be BMP independent, showed the most pronounced reductions in velocity length. While dI populations were not broadly affected, the dI1 trajectory was an exception, showing reduced velocity vectors in Smad1Δ and Smad5Δ cells compared to control (RA+BMP4 protocol, Fig. 7I, 7J). We sought to identify the genes driving velocity in dI1s and identified *Nav3* (neuronal navigator 3) and *Cadm2* (cell adhesion molecule 2) as the top-ranked genes with reduced velocity in both lines (Fig. 7K). Both genes have enriched expression in a subset of dI1s that are markedly lost in the Smad5Δ atlas (Fig. 7L). Nav3 (Martínez-López et al., 2005; van Haren et al., 2009) and Cadm2 (Frei et al., 2014) regulate cytoskeletal dynamics and cellular adhesion respectively, suggesting that Smad1Δ and Smad5Δ dI1s are impaired for neurite extension, axon guidance, and synapse formation.

While both Smad1Δ and Smad5Δ trajectories exhibit reduced RNA velocity, this effect is most pronounced for the Smad5Δ line. We thus propose Smad5 as the principal mediator of the transcriptional programs that maintain progenitor activity and regulate their progression into differentiated dI states (Fig. 7N).

## Discussion

In this study, we leveraged our stem cell model of the mouse dorsal spinal cord (Gupta et al., 2022) to investigate the roles of the R-Smad proteins mediating downstream BMP signaling during spinal cord development. Bulk and scRNA-seq datasets derived from control, Smad1Δ, and Smad5Δ mESC lines reveal that the R-Smads are required for distinct functions at multiple developmental decision points in the dI differentiation timeline. The loss of R-Smads does not prevent mESCs maintaining pluripotency (Fig. 2), or acquiring NMP identity (Fig. 3), suggesting that R-Smads are either dispensable or play redundant roles during the early stages of dI development. After these stages, their roles diverge, and Smad5 assumes major and reiterative roles. Smad5 is required [1] to bias NMPs towards a neural lineage, perhaps by preventing their differentiation into mesodermal derivatives (Fig. 5); [2] for the generation of dP1/dP2s and dI1/dI2s, which concomitantly limiting the specification of the intermediate dI4–dI6 populations (Fig.6); and [3] to maintain transcriptional dynamics across all dP populations (Fig. 7), given the observation that the rate of transcriptional change in dPs, measured by RNA velocity, is broadly dampened in the Smad5Δ line. By contrast, Smad1 acts to prevent progenitors adopting the RP fate, and restricts the size of dP1 pool.

These studies have additionally generated new resources, including ATAC-seq and RNA velocity atlases, that advance our understanding of the mechanisms underlying dorsal spinal cord development. These datasets also can be used to identify BMP4-responsive genes and genomic regions relevant to other developmental contexts where BMP signaling plays a regulatory role (Fig.4). Together, these findings begin to resolve how the Smad second messenger translates a common upstream signal into diverse cellular outcomes. Functional specificity is achieved, at least in part, through the divergent and non-redundant activities of individual downstream transducers.

### Model for Smad1/Smad5 function

The observation that Smad5 is the key mediator of the ability of BMP4 to drive dP1/dP2 specification and dI1/dI2 differentiation, is in general agreement with our previous studies *in vivo* (Hazen et al., 2012). These studies used conditional genetics to deplete either Smad1 or Smad5 in E10.5 mouse embryos. We found that the loss of Smad5, but not Smad1, resulted in decreased numbers of Atoh1 (Math1)^+^ dP1 progenitors and Lhx2^+^ dI1s. These studies *in vitro* go further, since we were able to dissect the requirement for the R-Smads at much finer resolution (Fig. 6I, 6L), as well as more globally across the differentiation behavior of all dPs (Fig. 7N) to give a more complete picture for how Smad5 vs Smad1 is required for dI differentiation. A additional major finding of our previous studies was that the loss of Smad1, but not Smad5, resulted in defects in axonal outgrowth (Hazen et al., 2012). While axon guidance phenotypes have not been the focus of these studies, we did observe that axon outgrowth is dramatically impaired in the Smad1Δ day 9 cultures (Fig. 6K) and that transcriptional signatures related to axonogenesis are downregulated (data not shown). Supporting this role of Smad1, Smad1 has been shown to be part of the regeneration-associated gene (RAG) response, a set of genes that are required for axon growth after injury (Saijilafu et al., 2013; Zou et al., 2009).

The opposing model posits that it is the combined strength of Smad1/5 activity, rather than the identity of the individual R-Smad, that dictates the outcome of the response to BMP signaling (Le Dreau et al., 2012; Le Dréau et al., 2014). Our findings raise two interpretative concerns about the concept of signal strength as a regulatory variable. First, what has been interpreted as a quantitative signal-strength effect may instead reflect a summation of qualitatively distinct contributions of Smad1 and Smad5 acting on different cellular processes, that are challenging to resolve *in vivo*. Second, we observe notably distinct results in our *in vitro* model system, under conditions of extrinsic BMP4 signaling, i.e., when BMP4 is supplemented directly in our differentiation protocol, compared to the intrinsic activation of BMP signaling in the RA protocol. It is only in the RA+BMP4 protocol that Smad1 and Smad5 appear to function redundantly in dI specification (Fig. 6D, 6E, 6G, 6H). We propose that elevating BMP4 concentrations, possibly to saturating levels, may mask the distinct functional contributions of individual R-Smads. This interpretation differs from the signal-strength model in an important respect: rather than signal strength dictating fate outcome with the R-Smads acting interchangeably, we propose that signal strength governs the extent to which the two R-Smads are functionally distinct. At high levels of BMP pathway activation which may be non-physiological, Smad1 and Smad5 function redundantly. In contrast, at reduced signal intensities, likely closer to physiological levels, the R-Smads exert qualitatively distinct effects on neural progenitor behavior.

### Mechanistic implications for BMP signal transduction

While Smad1 and Smad5 have previously suggested to be functionally redundant (Arnold et al., 2006; Retting et al., 2009; Wong et al., 2012), our work demonstrates that they can operate through distinct, non-overlapping mechanisms. This finding challenges the prevailing hypothesis that BMP pathway specificity arises from ligand–receptor combinations or extracellular modulators and instead highlights differential R-Smad usage as an underappreciated source of signaling diversity. Future studies will further determine the extent to which BMP pathway selectivity engages individual R-Smads in other tissues, and whether this phenomenon extends to Smad2 and Smad3, the R-Smads that mediate TGF-β signaling. Previous studies have suggested that Smad1 and Smad5 also have distinct roles in hematopoiesis (McReynolds et al., 2007).

While our findings suggest that Smad1 and Smad5 differentially regulate transcription, the mechanisms underlying these differences remain unresolved. R-Smads have intrinsically weak DNA-binding affinity, and their genomic engagement is largely stabilized through interactions with co-factors (Budi et al., 2017; Shi et al., 1998; Yoon et al., 2011). If Smad1 and Smad5 differed in their ability to recruit co-factors, their distinct transcriptional roles could result from altered DNA-binding affinity or site selectivity. The stoichiometry of R-Smad/co-Smad4 complexes could also influence DNA-binding affinity or the specific transcriptional output. Finally, phosphorylation and ubiquitination sites present in the proline-rich linker region connecting the R-Smad MH1 and MH2 domains have been shown to alter R-Smad nuclear import and export (Kamato et al., 2020; Wrighton et al., 2009) These sites differ between Smad1 and Smad5, which could differentially traffic the R-Smads to and from the nucleus, thereby contributing to their distinct functional roles.

### Non-canonical roles of Smad5

The studies also support prior observations that Smad5 has non-canonical functions, distinct from Smad1. Most notably, fluctuations in intracellular pH can drive non-activated Smad5 into the cytoplasm, where it binds hexokinase1 to enhance glycolysis and maintain cellular bioenergetic homeostasis (Fang et al., 2017). Our bioinformatic datasets consistently show that the loss of Smad5 results in more extensive transcriptomic disruption than loss of Smad1 (Fig. 4). After 24 hours of BMP4 treatment, 1054 genes are uniquely misregulated in the Smad5Δ atlas (Fig. 4P), compared to 175 in the Smad1Δ atlas (Fig. 4O). GO analysis support a role for Smad5 in cellular energetics with categories such as ‘mitochondrial ATP synthesis’ and “response to nutrient levels” in the downregulated genes (Fig. 4P). Given that metabolic programs are tightly coupled to neural progenitor proliferation and differentiation, this bioenergetic deficit may explain why the Smad5Δ progenitors show broadly decreased RNA velocities. Taken together, the gene sets of Smad5 regulated genes will be an invaluable resource to further examine additional non-canonical functions of Smad5.

### New insights into dI development from RNA velocity analyses

We performed an RNA velocity analysis to assess whether the R-Smads have differential impact on the transcriptional dynamics of dI development. This analysis resulted in several unexpected findings. First, we identified a transcriptionally distinct cluster, the multipotent dorsal progenitor (mdP) population, that the velocity vectors suggest gives rise to dP2–dP6 populations (Fig. 7A). This cluster could represent a long-hypothesized but elusive multipotent intermediate in dI development, that has the potential to give rise to restricted classes of dPs. This finding also suggests that dP1s arise independently of mdP, either earlier, or via a distinct lineage from the other dP subtypes. Second, BMP signaling, predominantly mediated by Smad5, is required to sustain the magnitude of the global velocity field during dP differentiation (Fig. 7G-7J). In the absence of Smad5, transcriptional dynamics are reduced across all dP clusters, including the subtypes previously considered to be BMP-independent (Fig. 7N) (Gupta and Butler, 2021). Thus, BMP signaling is a broader regulator of progenitor state transitions than previously appreciated. It both sustains the transcriptional programs that drive dPs through the differentiation process and directs patterning of the dorsal-most dIs.

Finally, we observed that, on differentiating, dIs generally show reduced transcriptional dynamics. There were two notable exceptions: the dI4 and early dI5 subtypes, which retained elevated velocity magnitudes (Fig. 7C). Notably, dI4s and dI5s are largest populations of interneurons in the mammalian spinal cord *in vivo* (Gupta et al., 2025; Roome et al., 2026) raising the possibility that their sustained transcriptional dynamics underlie this numerical expansion, allowing these populations to reach the sizes required to fulfill their functional roles in spinal circuits. Consistent with prolonged transcriptional activity, these populations diversify into a remarkable number of distinct functional subtypes over time (Gupta et al., 2025). Future studies will determine whether the elevated velocities observed in these lineages reflects ongoing fate refinement, continued proliferative activity, or delayed terminal differentiation.

### New resources to identify the key genes regulating dorsal spinal cord development

The mechanisms that pattern the dorsal spinal cord and thereby specify distinct dP and dI populations remain incompletely defined. These studies have also developed several resources, that can be used to identify the genes that regulate BMP signaling and drive dP/dI specification.

We generated an ATAC-Seq atlas mapping the chromatin accessibility changes resulting from BMP4 treatment, together with matched bulk RNA-Seq data from control, Smad1Δ and Smad5Δ lines. Together these resources permit BMP-dependent chromatin remodeling to be correlated directly to transcriptional responses. Using this approach, we identified *Wfikkn1* as a BMP4-induced gene in the developing dorsal spinal cord. Wfikkn1 is a secreted protein that can bind to BMP2 and BMP4, but sequesters, rather than antagonizes, them (Szlama et al., 2010). Thus, Wfikkn1 may function to fine-tune BMP4 signaling, rather than acting as a classical antagonist. In this model, Wfikkn1 functions in a negative-feedback loop; it is induced by BMP4 to sequester BMP4, and thereby shape the spatial range of BMP signaling during dorsal spinal cord development. This model parallels FGF and Hedgehog signaling which similarly induce their own attenuators, Sprouty and Patched respectively. Future studies will assess the role of *Wfikkn1* in the developing spinal cord *in vivo*, including determining its spatial location.

We have also generated an RNA velocity atlas, which can be used to identify genes with dynamic transcription patterns in specific dP populations. These dP-specific candidate driver genes may play essential roles enabling progenitors to transition into their respective dP subtypes, thereby addressing a key gap in our understanding of dorsal spinal cord development. Defining these drivers has translational potential: the current directed-differentiation protocols do not produce pure dI subtypes, and under-represent some lineages. By clarifying the transcriptional programs and developmental trajectories that guide dP progression, these genes could be used to refine existing differentiation strategies and generate defined dI populations for regenerative or cell-replacement applications.

### Summary

Taken together, these findings suggest that Smad1 and Smad5 have unique roles in the dorsal spinal cord, rather than being functionally interchangeable. We have also developed key resources that will help enable future studies to uncover the genetic mechanisms that give rise to these differences.

## Competing interests

There are no competing interests to declare.

## Author Contributions

S.G._1_, S.G._2_ and S.J.B. conceived the project. S.G._2_ designed and led the CRISPR experiments to generate Smad1Δ and Smad5Δ mESC lines and was assisted by B.C., and A.D. S.G._1_ performed all *in vitro* differentiation experiments, with G.G.D.R. and C.R. assisting with cell culture maintenance and sample collection. S.G._1_ analyzed the bulk and scRNA-seq data while Y.V. performed the ATAC-seq analysis. S.G._1_ performed the IHC analyses and C.R. performed the Western analyses. This paper was written by S.G_1_., and S.J.B. and edited by S.G_1_., S.G._2_ and S.J.B.

## Acknowledgements

We would like to thank Aparna Bhaduri, Bill Lowry, Karen Lyons, Bennett Novitch and members of the Butler and Novitch laboratories for discussions. These studies were supported by a Whitcome pre-doctoral fellowship and a fellowship from the National Institutes of Health (NIH) Training Grant in Genomic Analysis and Interpretation (T32HG002536) to S.G._1_, a UCLA Broad Stem Cell Research Center (BSCRC) postdoctoral fellowship to S.G._2_, a Lawrence Kruger Undergraduate Research Scholarship to A.D, the California Institute for Regenerative Medicine (CIRM) Bridges to Research program (EDUC2-12718) to B.C., C.R., G. G. del R., and Y.V., and grants to S.J.B. from the NIH (R01NS123187) and the Marcus Foundation (4492).

## Notes

### Competing Interest Statement

The authors have declared no competing interest.

